# A multidimensional benchmark of bulk RNA-seq simulators for task-specific evaluation of simulation fidelity

**DOI:** 10.64898/2026.11.25.768677

**Authors:** Nazia Riasat, Megan Orr

## Abstract

Bulk RNA-seq simulators are widely used to develop and evaluate statistical methods, but their suitability depends on whether they preserve the properties of the data required for the intended downstream analysis. We benchmarked seven widely used simulators across five real bulk RNA-seq reference datasets using a unified framework that evaluated five complementary dimensions of simulation fidelity: marginal distribution fidelity, dependence structure, co-expression network topology, stability across repeated simulations, and differential-expression recovery. Simulator performance varied substantially across these categories, and no method consistently reproduced all properties of the reference data. Rankings differed across categories, with only moderate agreement between evaluation objectives, indicating that these measures capture distinct dimensions of simulation fidelity rather than a single underlying notion of performance. Under the network construction criteria used here, network topology was the most difficult property to reproduce consistently across simulators, whereas individual methods showed strengths in marginal distributions, dependence structure, repeated-run stability, or differential-expression recovery. Although an aggregate score provides a concise descriptive summary, it can hide meaningful differences between evaluation objectives. These results show that bulk RNA-seq simulation fidelity is inherently multidimensional and support a task-specific framework for simulator evaluation and selection, in which simulator choice is guided by the statistical and biological properties required by the downstream analysis rather than by a single overall ranking.

## 1 Introduction

Bulk RNA sequencing remains widely used to quantify gene expression across biological conditions and provides the basis for differential-expression testing, pathway analysis, clustering, and co-expression network analysis Wang et al. [2009], Love et al. [2014], Law et al. [2014], Langfelder and Horvath [2008]. These analyses do not depend on the same properties of an expression dataset. Differential-expression methods are influenced by count distributions and mean–variance relationships, whereas clustering, dimension reduction, and network analyses depend more strongly on relationships among samples and genes Love et al. [2014], Law et al. [2014], Ballouz et al. [2015]. Reproducing the marginal distribution of RNA-seq counts is therefore not enough to establish that a simulated dataset also reproduces its multivariate or network structure.

Simulation is widely used to evaluate statistical methods because it provides controlled settings in which sample size, sequencing depth, group structure, and differential-expression status can be specified. Its usefulness depends on whether the simulated data preserve the properties needed for the analysis being evaluated. RNA-seq simulation fidelity therefore involves several aspects of the data rather than a single summary measure. Previous benchmarking studies, including work in single-cell settings, have also shown that simulator performance depends on the criterion being assessed and that the choice of simulator can influence conclusions about downstream methods Soneson and Robinson [2018], Cao et al. [2021], Crowell et al. [2023], Shakola et al. [2022].

Bulk RNA-seq simulators were developed for different purposes and use different strategies to generate expression data. compcodeR and powsimR use model-based count generation and were developed mainly for differential-expression benchmarking and power analysis [Soneson, 2014, Vieth et al., 2017]. SimSeq instead uses observed RNA-seq data and does not require a specific parametric count model [Benidt and Nettleton, 2015]. seqgendiff also starts from observed counts and introduces controlled differential-expression signals through binomial thinning, allowing much of the variation in the original data to remain [Gerard, 2020]. SPsimSeq uses a semi-parametric approach designed to reproduce gene-level expression distributions while retaining correlation patterns observed in the reference data Assefa et al. [2020]. Splatter was developed as a flexible framework for simulating single-cell RNA-seq data under different biological and experimental settings Zappia et al. [2017], whereas SimBu generates pseudo-bulk expression profiles from single-cell data and allows the underlying cell-type composition to be specified [Dietrich et al., 2022]. These differences in purpose and construction can affect which features of a reference dataset are reproduced well.

Several studies have compared RNA-seq simulation methods, although the properties examined have varied. For bulk RNA-seq, previous comparisons have considered distributional similarity, differential-expression recovery, correlation structure, and class separation [Soneson and Robinson, 2018, Shakola et al., 2022]. Gerard [2020] also examined different simulation strategies and showed that conclusions from method evaluations can depend on how the simulated data are generated. Broader comparisons have been carried out for single-cell RNA-seq. Zappia et al. [2017] compared several existing approaches when introducing Splatter, Cao et al. [2021] evaluated 12 simulators across 35 datasets, and Crowell et al. [2023] later compared 16 methods using both data characteristics and downstream analyses. Together, these studies show that simulator performance depends on which aspect of the data is being evaluated. However, bulk RNA-seq simulators have been less extensively compared across marginal distributions, gene-gene and sample-sample dependence, co-expression network topology, repeated-run stability, and differential-expression recovery within the same set of reference datasets.

We compare seven RNA-seq simulators across five real bulk RNA-seq reference datasets using a common evaluation framework. Simulation quality is examined across five categories: marginal distribution fidelity, dependence and multivariate structure, co-expression network topology, stability across repeated simulations, and differential-expression recovery. Output availability is tracked separately so that failed or non-evaluable simulations are not treated as successful observations. We also compare category-specific performance profiles to assess whether the evaluation dimensions provide similar or distinct information.

The framework is intended to support simulator selection according to the properties required by the intended downstream analysis rather than by a single overall measure (Fig. 1).

**Figure 1:**
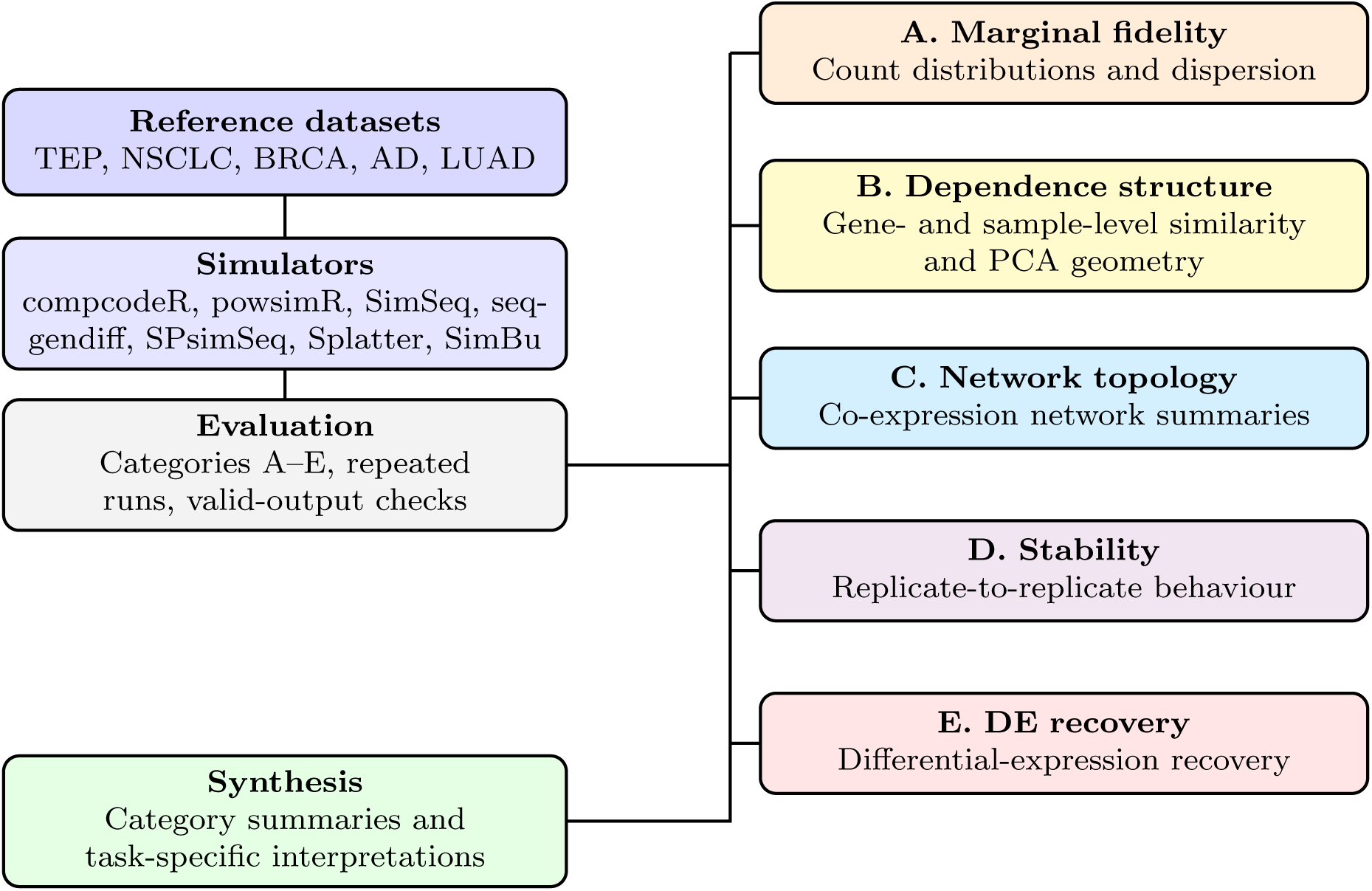
Benchmark design. Five real bulk RNA-seq datasets were used as references for seven simulation methods. Simulated datasets were evaluated across five complementary dimensions: marginal distribution fidelity (A), dependence and multivariate structure (B), co-expression network topology (C), stability across repeated simulations (D), and differential-expression recovery (E). Category-level results were then compared to characterize task-specific simulator performance.

## 2 Methods

### 2.1 Reference datasets and preprocessing

Five publicly available bulk RNA-seq datasets were used as references: tumor-educated platelets (TEP; *n* = 100), non-small-cell lung cancer (NSCLC; *n* = 25), breast cancer (BRCA; *n* = 200), Alzheimer’s disease (AD; *n* = 150), and lung adenocarcinoma (LUAD; *n* = 100) [Best et al., 2015, National Center for Biotechnology Information, 2015, 2023, The Cancer Genome Atlas Network, 2012, The Cancer Genome Atlas Research Network, 2014, Wilks et al., 2021]. The selected cohorts contained two biological groups in each dataset and differed in sample size, sequencing depth, and count sparsity. Dataset-specific sample selection and filtering rules are given in Supplementary Section A1.

After samples with missing or ambiguous group labels had been removed, the count matrices were subjected to the common benchmark filter. TMM normalization [Robinson and Oshlack, 2010] and log-counts per million (logCPM) transformation were used only to rank genes by between-sample variance; the underlying raw counts were retained for simulator fitting and count-level evaluation. The 3000 genes with the largest logCPM variance were retained for the primary count-level benchmark so that all simulators were compared at the same gene dimension. For co-expression network analyses, the top 1000 genes from the same variance ranking were used consistently for the reference and simulated datasets. Gene-set-size sensitivity for the marginal and dependence analyses was examined in BRCA using 3000, 5000, 10000, and 15000 genes.

### 2.2 Simulation design and output handling

Seven simulators were evaluated: compcodeR, powsimR, SimSeq, seqgendiff, SPsimSeq, Splatter, and SimBu. These methods represent model-based, non-parametric, semi-parametric, empirical-reference, single-cell, and pseudo-bulk simulation strategies [Soneson, 2014, Vieth et al., 2017, Benidt and Nettleton, 2015, Gerard, 2020, Assefa et al., 2020, Zappia et al., 2017, Dietrich et al., 2022]. Package versions, simulator-specific parameters, random-number settings, and implementation details are reported in Supplementary Section A11. Each simulator was calibrated separately to each reference dataset for which evaluable output could be produced. Simulated matrices were generated to match the reference gene dimension and, where supported by the simulator, the reference sample size and two-group structure. Ten independent simulation replicates were generated for each evaluable simulator–dataset combination.

For Categories A, B, C, and E, replicate-level metric values were summarized by their mean before scoring. Replicate-to-replicate variation was evaluated separately as Category D. Simulation failures and undefined metrics were recorded separately. If a simulator did not produce a usable count matrix for a dataset, that run was marked as unavailable. If the simulated data were valid but a downstream metric could not be calculated, the metric was recorded as undefined. These cases were not assigned a performance score, and coverage was reported separately from fidelity. Differential-expression recovery was also left unscored when simulated group labels could not be matched reliably to the reference groups. Full coverage information is provided in Supplementary Section A6.

### 2.3 Evaluation categories

Simulation quality was evaluated across five categories chosen to represent distinct properties of bulk RNA-seq data.

#### Category A: marginal distribution fidelity

Agreement in marginal count behaviour was evaluated using several distributional summaries. The mean–variance relationship across genes was compared using integrated squared error (ISE) [Wand and Jones, 1995]. Kolmogorov–Smirnov (KS) distances were used to compare the distributions of logCPM values, library sizes, the number of zero counts per gene, the number of zero counts per sample, and the biological coefficient of variation (BCV) [Massey, 1951, Smirnov, 1948]. For all Category A metrics, smaller values indicate closer agreement between the simulated and reference data.

#### Category B: dependence and multivariate structure

Dependence preservation was evaluated from several complementary perspectives. Frobenius distance was used to measure the overall difference between real and simulated gene–gene and sample–sample relationship matrices. The RV coefficient [Robert and Escoufier, 1976] measured similarity between the corresponding multivariate dependence structures, while Procrustes *M* ^2^ quantified differences in their geometric configuration [Gower, 1975]. Similarity in principal-component structure was assessed by correlating the cumulative proportions of variance explained across principal components. Smaller Frobenius distances and Procrustes *M* ^2^ values indicate better agreement, whereas larger RV coefficients and variance-explained correlations indicate better agreement.

#### Category C: co-expression network topology

Network agreement was assessed using the relative error between simulated and reference networks for four topology summaries: density, modularity [Newman, 2006], average degree, and clustering coefficient [Watts and Strogatz, 1998]. Smaller relative errors indicate closer agreement with the reference network. A network-usability check was applied before these quantities were scored because relative topology summaries are not meaningful for degenerate or nearly empty graphs. AD was excluded from the primary Category C evaluation because its reference network did not satisfy the pre-specified network-usability criterion. For the remaining datasets, simulator outputs that failed the usable-network criterion were assigned the least favourable Category C score of 1 for that dataset. Network construction, the us-ability criterion, relative-error calculation, and sensitivity analyses are described in Supplementary Sections A2 and A6.

#### Category D: stability across repeated simulations

Replicate-to-replicate variation was evaluated separately as Category D because stability reflects reproducibility of simulator output rather than agreement with a reference-data property. For each simulator–dataset combination, replicate-to-replicate variation was calculated for the replicate-based evaluation metrics from Categories A, B, C, and E. For each replicate-based metric *x*, variation was summarized using the coefficient of variation [Reed et al., 2002],

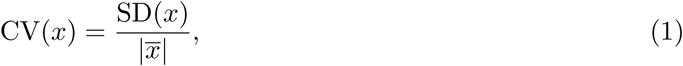

when the metric was defined and its mean was non-zero. The available metric-level CVs were then combined within each simulator–dataset combination to obtain an overall stability summary. These summaries were rank-normalized across simulators, with lower values indicating less run-to-run variation. Category D was therefore interpreted as a measure of simulation reproducibility rather than biological realism. Details of the metric-level CV calculation, aggregation, and treatment of zero or undefined quantities are provided in Supplementary Section A2.

#### Category E: differential-expression recovery

Differential-expression analysis was applied to the reference and simulated datasets using matched group labels where available. Recall and Jaccard [Jaccard, 1901] overlap between reference and simulated differential-expression sets were used as the primary Category E summaries. ROC AUC, precision–recall AUC [Saito and Rehmsmeier, 2015], precision, F1 score, and type I error were retained as supporting diagnostics. Larger values indicate better recovery for recall, Jaccard overlap, ROC AUC, precision–recall AUC, precision, and F1; smaller values indicate better control for type I error. Complete metric definitions are provided in Supplementary Section A2.

### 2.4 Rank-normalized scoring and aggregation

The evaluation metrics have different units and numerical ranges, so raw values were not averaged directly. For each dataset and metric, simulators with valid results were ranked in the appropriate direction. If *r_m,k_* denotes the rank of simulator *m* for metric *k* and *N_k_* is the number of simulators with valid values for that dataset–metric combination, the rank was mapped to

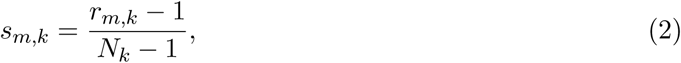

so that *s_m,k_*= 0 represents the strongest valid result and *s_m,k_* = 1 the weakest. Tied raw metric values received the corresponding tied rank before normalization.

Within Categories A, B, C, and E, normalized scores for the available primary scoring metrics were averaged to obtain a dataset-level category score. Final category scores for A, B, C, and E were obtained by taking the median of the corresponding dataset-level scores across evaluable reference datasets. Category D was constructed separately from the replicate-variation analysis described above; its final pooled score was the arithmetic mean of the five corrected dataset-level stability scores. After accounting for metric direction, lower normalized scores indicate stronger performance in every reported category.

Because the five evaluation categories quantify complementary properties of simulation fidelity, no single category was considered sufficient to characterize simulator performance. For completeness, an equal-weight five-category composite score was computed as a secondary descriptive summary using

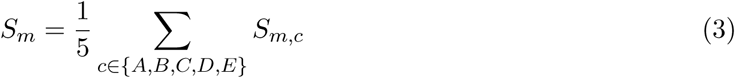

where *S_m,c_* is the pooled score for simulator *m* in category *c*. Because this composite gives equal weight to conceptually different evaluation targets, category-specific performance profiles formed the primary basis for interpretation, whereas the composite score was used only as a descriptive overview. Metric coverage, defined as the proportion of expected dataset–metric combinations with valid values, was reported alongside performance and was not folded into the composite score except for the explicit Category C usable-network penalty.

### 2.5 Uncertainty, sensitivity, and supporting analyses

Sensitivity of the composite summary to reference-dataset composition was assessed by resampling the five reference datasets with replacement for 2000 bootstrap iterations and recomputing the composite summary and corresponding simulator ordering. Median bootstrap rank and the 2.5th and 97.5th percentiles of the bootstrap rank distribution were reported. These intervals quantify sensitivity to the composition of the reference-dataset set rather than uncertainty in an underlying population parameter. Sensitivity to gene-set size was evaluated in BRCA at 3000, 5000, 10000, and 15000 genes. Sample-size analyses were treated as supporting experiments rather than as an additional benchmark category. compcodeR and Splatter were evaluated under a common external differential-expression workflow over increasing sample sizes, whereas powsimR was examined with its package-specific power procedure. Full settings are provided in Supplementary Section A13. Functional coherence of simulated co-expression networks was assessed separately using EGAD neighbour voting [Ballouz et al., 2017] with MSigDB Hallmark gene sets [Liberzon et al., 2015]. This analysis was applied across the benchmark datasets and simulators as a biological check on the network results and was not included in the Category C score. Methods and results are provided in Supplementary Section A12.

### 2.6 Statistical comparisons and reproducibility

Agreement among category-specific simulator performance profiles was quantified within each dataset using Kendall’s *τ* [Kendall, 1938]. Category D was excluded from this comparison because it measures repeated-run stability rather than fidelity to a feature of the reference data. We also compared simulators pairwise to assess whether the observed differences in individual metrics were consistent across repeated simulations and reference datasets. Within each dataset and metric, replicate values from two simulators were compared using the Wilcoxon rank-sum test, which does not require a normal distribution. Because many simulator pairs, metrics, and datasets were tested, the resulting *p*-values were adjusted using the Benjamini–Hochberg procedure to limit the false discovery rate. Cliff’s delta was reported alongside the *p*-values to describe both the size and direction of the differences. We also used paired Wilcoxon signed-rank tests to compare matched simulator summaries across reference datasets when both simulators had valid results. Because only five reference datasets were available for the across-dataset comparisons, these analyses were treated as supporting evidence and were not used to determine category scores or the overall simulator ranking. Full results are reported in Supplementary Section A9.

All analyses were performed in R [R Core Team, 2024] using a reproducible pipeline covering dataset preprocessing, simulator calibration, repeated simulation, metric calculation, scoring, ro-bustness analyses, and figure generation. Software versions and simulator settings are provided in Supplementary Section A11. Consequently, recommendations throughout this benchmark are based primarily on category-specific performance rather than the composite summary.

## 3 Results

### 3.1 Simulation performance differs across evaluation dimensions

Simulation performance depended strongly on the property being evaluated (Figure 2). Different simulators performed best for different features of the RNA-seq data. seqgendiff performed best overall for marginal count fidelity and dependence structure, SimSeq for co-expression network topology among methods producing usable networks, and compcodeR for repeated-run stability. seqgendiff and SimSeq performed equally well on the pooled differential-expression recovery score, although their dataset coverage differed substantially (Table 1). These results indicate that simulator performance is multidimensional rather than well represented by a single criterion.

**Figure 2:**
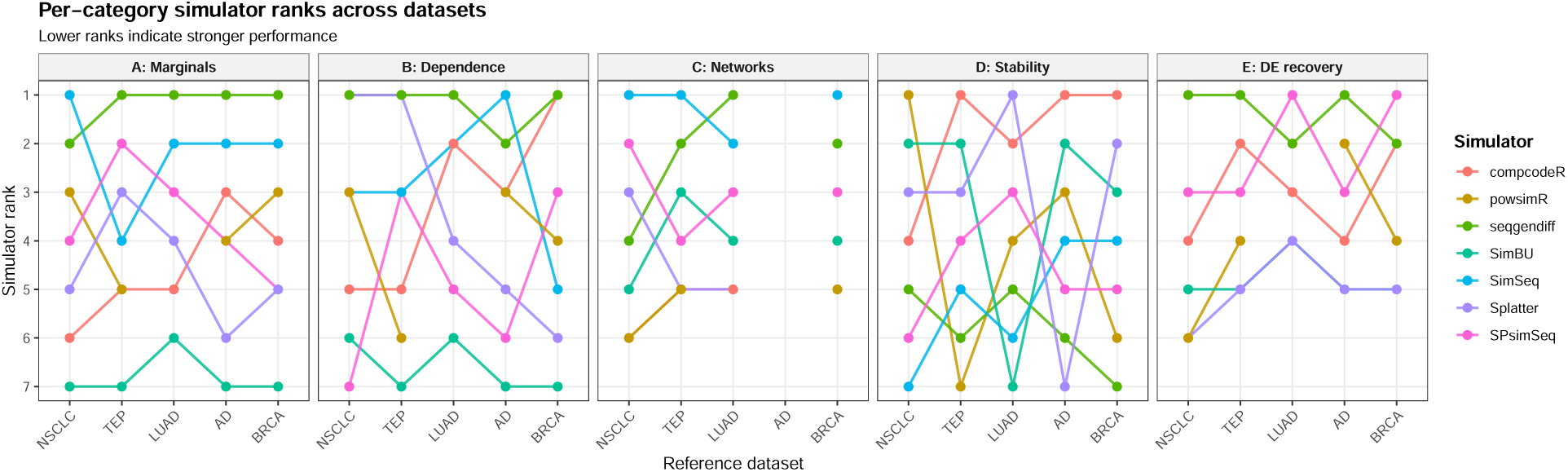
Dataset-specific simulator performance across the five evaluation categories. Rank 1 denotes the lowest category score within a dataset. Lines connect results for the same simulator across datasets; unavailable results are left unconnected. Rankings changed with both the evaluation target and the reference dataset.

**Figure 3:**
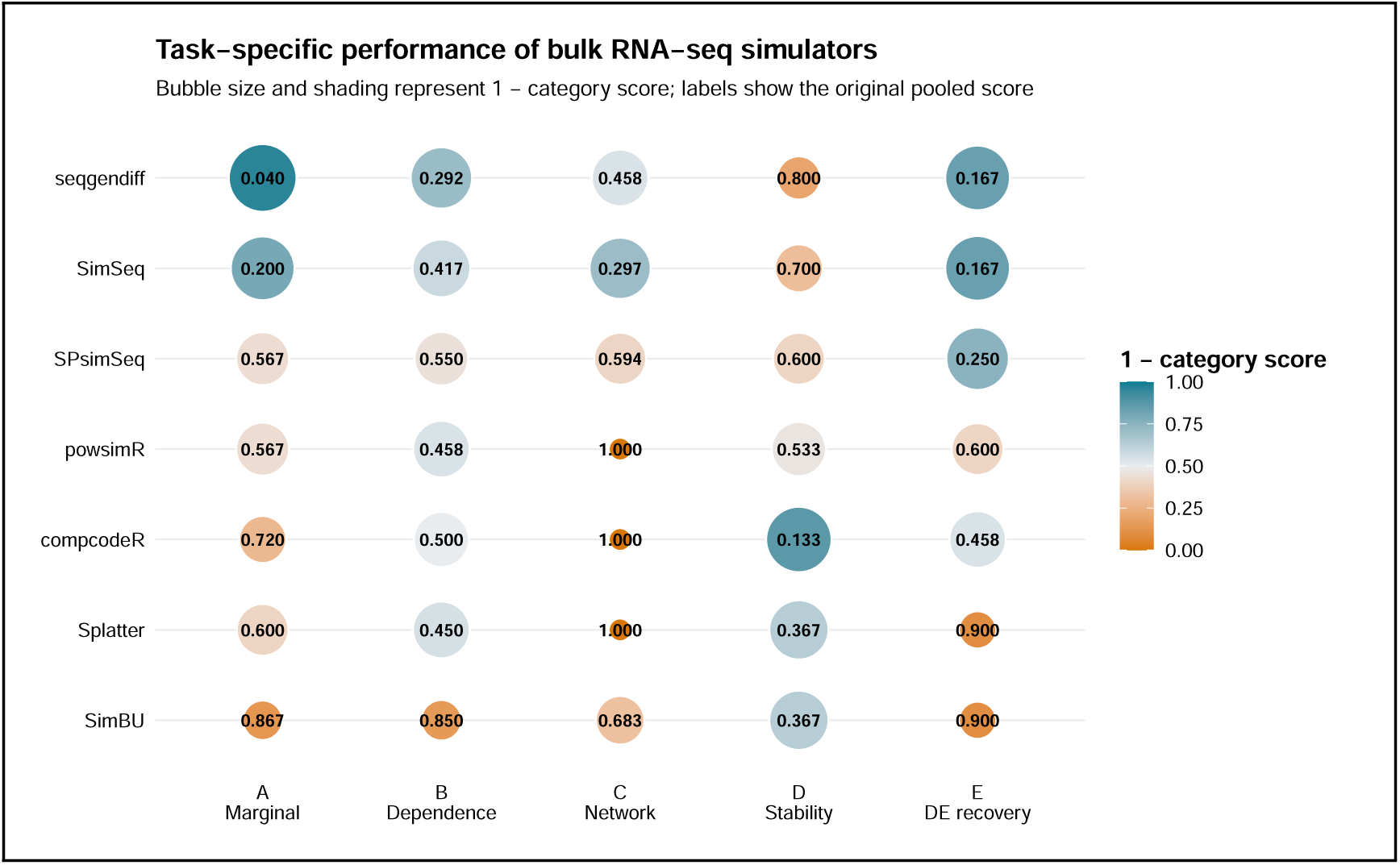
Pooled performance of the seven bulk RNA-seq simulators across the five evaluation categories. Numbers inside the circles are pooled rank-normalized category scores, with lower values indicating stronger performance within the corresponding evaluation dimension. Circle size and shading represent 1 − score, so larger circles indicate lower category scores. Category D summarizes stability across repeated simulations, whereas Categories A, B, C and E summarize agreement with the reference data or downstream recovery.

**Table 1:**
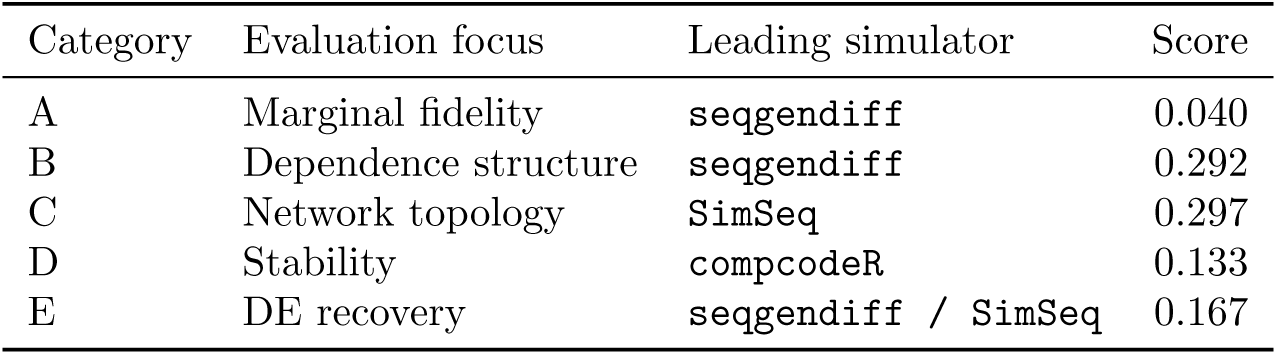
Leading simulator in each evaluation category based on pooled rank-normalized scores. Lower values indicate stronger performance. Category C includes the usable-network rule for degenerate or nearly empty networks and excludes AD because the reference network was not evaluable. Category D measures repeated-run stability rather than agreement with a feature of the reference data.

| Category | Evaluation focus | Leading simulator | Score |
|---|---|---|---|
| A | Marginal fidelity | seqgendiff | 0.040 |
| B | Dependence structure | seqgendiff | 0.292 |
| C | Network topology | SimSeq | 0.297 |
| D | Stability | compcodeR | 0.133 |
| E | DE recovery | seqgendiff / SimSeq | 0.167 |

The category rankings were only partly consistent. Marginal fidelity and dependence showed the greatest agreement, with a median Kendall’s *τ* = 0.552 across datasets. Dependence and network topology showed the lowest agreement, with a median *τ* = 0.080 across the four datasets for which Category C was evaluable. This low agreement is plausible because the two categories capture different aspects of structure: dependence metrics summarize broader relationships among genes and samples, whereas network topology depends on how those relationships translate into edges and higher-order graph properties. Category D was excluded from this comparison because it measures repeated-run variation rather than agreement with a reference-data property. Full pairwise results are reported in Supplementary Section A7.

### 3.2 Marginal count fidelity is strongest for reference-based simulations

Category A evaluated agreement with the marginal distribution of RNA-seq counts. For Category A, seqgendiff performed best overall in Category A (lowest pooled score 0.040), followed by SimSeq (0.200). The remaining scores ranged from 0.567 for SPsimSeq and powsimR to 0.867 for SimBu; the powsimR summary was based on four datasets because evaluable LUAD output was not available.

The dataset-level pattern was similar. seqgendiff had the highest Category A performance for TEP, BRCA, AD and LUAD, whereas SimSeq had the highest performance for NSCLC. At the metric level, seqgendiff had the most favourable pooled value for each of the six marginal summaries: mean–variance ISE, logCPM distribution, library size, zeros per gene, zeros per sample and biological coefficient of variation. The size of the discrepancy nevertheless varied by metric and dataset, so agreement in one marginal property did not imply comparable agreement in all others. Metric-level and per-dataset results are given in Supplementary Sections A4 and A5.

### 3.3 Dependence preservation varies across metrics

Category B evaluated preservation of dependence and multivariate structure. Category B showed a less uniform pattern than Category A. seqgendiff performed best overall in Category B (lowest pooled score 0.292), followed by SimSeq (0.417); the remaining scores ranged from 0.450 for Splatter to 0.850 for SimBu. The powsimR summary again reflected four-dataset coverage because LUAD was unavailable.

No single simulator led every dependence metric. seqgendiff had the highest performance in gene–gene Frobenius distance and PVE correlation, SimSeq had the highest performance in sample–sample Frobenius distance, powsimR had the highest performance in RV coefficient, and compcodeR had the highest performance in Procrustes *M* ^2^. At the dataset level, seqgendiff had the lowest or tied-lowest Category B score in TEP, NSCLC, BRCA and LUAD indicating highest performance across these datasets, whereas SimSeq had the highest performance in AD. Thus, gene-level dependence, sample-level structure and PCA geometry did not yield the same simulator ordering. Full results are provided in Supplementary Sections A4 and A5.

### 3.4 Network evaluability separates the simulators before topology is compared

Category C first required an interpretable co-expression network. compcodeR and powsimR produced near-zero-density networks on each evaluable reference dataset, while Splatter did so on three of four. These simulator–dataset combinations met the predefined degeneracy criterion and were assigned the least favourable Category C score of 1 for the affected dataset. In contrast, seqgendiff, SimSeq, SPsimSeq, and SimBu produced usable networks across BRCA, LUAD, NSCLC, and TEP. AD was omitted from the primary topology comparison because its reference network did not meet the evaluability criterion. After excluding AD, SimSeq performed best overall in Category C (lowest pooled score 0.297), followed by seqgendiff (0.458), SPsimSeq (0.594), and SimBu (0.683). compcodeR, powsimR, and Splatter received pooled scores of 1.000 because their simulated networks failed the usability rule on most or all evaluable datasets. The metric-level ordering was less uniform: seqgendiff was closer for several global network summaries, whereas SimSeq and SPsimSeq showed stronger agreement for clustering. SimBu passed the network-usability criterion but produced substantially denser networks than the corresponding references. At the dataset level, SimSeq had the highest performance in Category C for BRCA, NSCLC, and TEP, whereas seqgendiff had the best score for LUAD. Functional coherence was assessed separately using neighbour-voting prediction of MSigDB Hallmark annotations and was not included in Category C scoring. These results are reported in Section A12; network validity and metric coverage are reported in Supplementary Section A6.

### 3.5 Repeated-run stability is distinct from fidelity

Category D produced a different ordering from the fidelity categories. compcodeR showed the strongest repeated-run stability (lowest pooled stability score 0.133), followed by Splatter and SimBu (0.367 each), powsimR (0.533), SPsimSeq (0.600), SimSeq (0.700) and seqgendiff (0.800). Lower values indicate less replicate-to-replicate variation. The dataset-level leaders also differed: compcodeR ranked highest for AD, BRCA and TEP, powsimR for NSCLC, and Splatter for LUAD. Thus, the simulator with the strongest marginal and dependence scores, seqgendiff, was the least stable in the pooled Category D summary, while compcodeR was the most stable. These scores describe reproducibility of repeated output and not closeness to the reference data.

### 3.6 Differential-expression recovery differs from the other category rankings

Category E used recall and Jaccard overlap as the primary recovery summaries, with ROC AUC, PR AUC, precision, F1 score and type I error retained as supporting diagnostics. seqgendiff and SimSeq tied for the strongest pooled Category E performance (shared the lowest pooled score 0.167), followed by SPsimSeq (0.250), compcodeR (0.458), powsimR (0.600), and SimBu and Splatter (0.900). The tie between seqgendiff and SimSeq reflected very different coverage. seqgendiff was evaluable across all five reference datasets, whereas the SimSeq Category E score was based on NSCLC only. At the individual-metric level, SPsimSeq had the highest pooled recall, while compcodeR had the highest pooled Jaccard overlap and led several supporting DE summaries. Dataset-level leaders also changed with the reference: seqgendiff had the lowest Category E score for TEP and AD and shared the lowest score on NSCLC with SimSeq, whereas SPsimSeq had the lowest score for BRCA and LUAD.

The differences across datasets may reflect variation in sample size, expression variability, dispersion, heterogeneity, and the strength of the differential-expression signal in the reference data. A simulator may therefore reproduce one dataset well but behave differently under another reference setting. This suggests that performance is shaped by both the simulation method and the characteristics of the reference dataset, rather than being a fixed property of the simulator. Thus, downstream DE recovery could not be inferred from marginal, dependence, or network performance alone.

### 3.7 Category-specific results give different task summaries

The five categories describe complementary aspects of simulator behaviour. Marginal fidelity, dependence structure, network topology, repeated-run stability, and differential-expression recovery produced distinct performance profiles across simulators. Rather than supporting a universal ordering, the benchmark indicates that simulator selection should follow the statistical property most relevant to the downstream application. Table 2 summarizes the method with the lowest pooled score for each target without treating the result as a universal ordering. Marginal fidelity and dependence both favoured seqgendiff; network topology favoured SimSeq; repeated-run stability favoured compcodeR; and differential-expression recovery produced a pooled tie between seqgendiff and SimSeq, with substantially broader coverage for seqgendiff.

**Table 2:** Task-specific simulator recommendations based on evaluation objective. The table reports the best-performing simulator for each evaluation target rather than a single overall choice.

| Evaluation target | Main evidence | Best-performing simulator | Interpretation |
|---|---|---|---|
| Marginal count fidelity | Category A marginal summaries | seqgendiff | Lowest pooled Category A score. |
| Dependence and multivariate structure | Category B dependence summaries | seqgendiff | Lowest aggregate score, although individual metrics favoured different methods. |
| Co-expression network topology | Category C topology summaries | SimSeq | Lowest reported pooled topology score among methods producing usable networks throughout. |
| Repeated-run stability | Category D replicate variation | compcodeR | Lowest pooled replicate-variation score; stability is separate from fidelity. |
| Differential-expression recovery | Category E recovery summaries | seqgendiff / SimSeq | Same pooled score, but seqgendiff had substantially broader dataset coverage. |

### 3.8 Composite summaries are sensitive to reference-dataset composition

With equal weight assigned to the five evaluation categories, the final composite scores were 0.351 for seqgendiff, 0.356 for SimSeq, 0.512 for SPsimSeq, 0.562 for compcodeR, 0.632 for powsimR, 0.663 for Splatter, and 0.733 for SimBu. The difference between the first two composite scores was only 0.005. Their category profiles were nevertheless different: seqgendiff had lower marginal and dependence scores, whereas SimSeq had the lower network-topology score, and compcodeR had the lowest stability score. Sensitivity of the aggregate ordering to reference-dataset composition was assessed using 2000 bootstrap resamples of the five datasets. The median bootstrap rank was 1 for seqgendiff, with a 2.5th–97.5th percentile rank interval of 1–2, and 2 for SimSeq, also with an interval of 1–2. SPsimSeq had median rank 3 with an interval of 2–4. The corresponding intervals were 2–7 for compcodeR, 3–7 for powsimR, 3–7 for Splatter, and 5–7 for SimBu. The overlap between the two leading methods and the broader intervals among the middle-ranked methods show that the aggregate ordering is sensitive to which reference datasets contribute to the summary. The composite is therefore retained as a descriptive overview rather than as a substitute for the category-specific results. Figure 4 summarizes the pooled scores and bootstrap rank intervals, with full bootstrap results reported in Supplementary Section A8.

**Figure 4:**
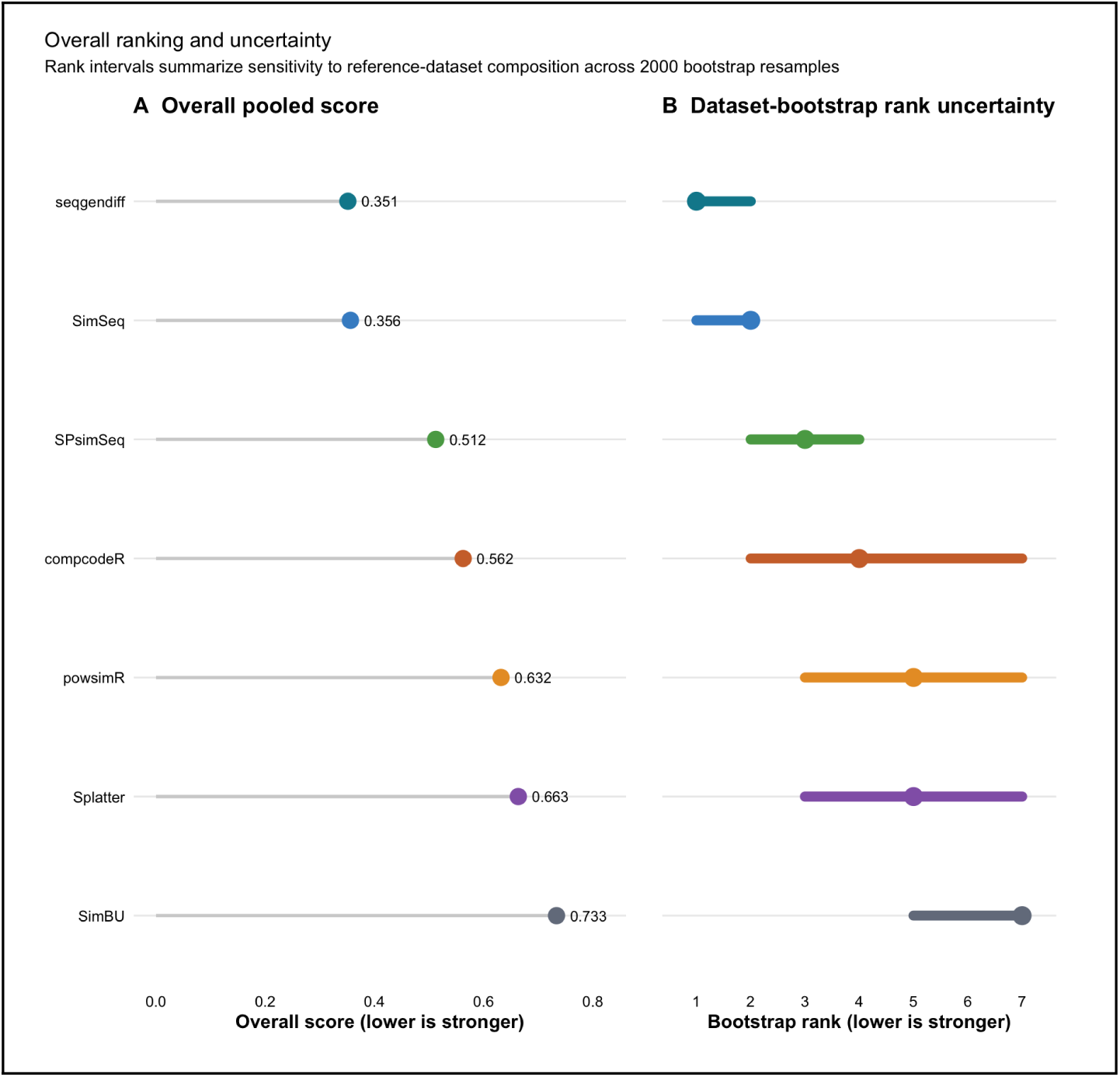
Overall simulator ranking and sensitivity to reference-dataset composition. A: Composite score obtained by assigning equal weight to the five evaluation categories; lower values indicate stronger aggregate performance. B: Median simulator rank and 2.5th–97.5th percentile rank intervals from 2000 bootstrap resamples of the five reference datasets. The intervals describe sensitivity of the aggregate ordering to dataset composition rather than uncertainty in a population parameter.

## 4 Discussion

This benchmark shows that bulk RNA-seq simulator performance is multidimensional. Across seven simulators and five reference datasets, different methods reproduced different properties of the reference data, including marginal count behaviour, dependence structure, co-expression network topology, and differential-expression recovery, while repeated-run stability captured a separate aspect of simulator reproducibility. No simulator performed uniformly across these evaluation targets. Instead, performance depended on the objective being considered, indicating that simulator selection should be guided by the statistical and biological properties required for the intended downstream analysis rather than by a single overall measure. The results therefore support a task-specific view of RNA-seq simulation and show that the evaluation categories provide complementary rather than redundant information [Soneson and Robinson, 2018, Shakola et al., 2022, Cao et al., 2021].

The contrast between marginal and multivariate behaviour illustrates this principle clearly. seqgendiff reproduced the marginal summaries closely and also achieved the strongest overall dependence profile, but no simulator consistently performed best across every dependence metric. Gene–gene correlation structure, sample–sample geometry, principal-component variance, and Procrustes alignment each captured different characteristics of multivariate structure. These findings emphasize that agreement in gene-wise count distributions does not mathematically imply agreement in their joint distribution. Two simulated datasets may reproduce similar means, dispersions, and sparsity while differing substantially in gene correlations or sample relationships. Consequently, dependence should be viewed as a collection of complementary properties rather than a single characteristic of simulated RNA-seq data.

The observed patterns are broadly consistent with how the simulators were designed, although the benchmark does not identify which specific design choices caused the differences in performance. seqgendiff and SimSeq retain substantial information from observed expression profiles, whereas compcodeR, powsimR, and Splatter rely more on fitted probabilistic models [Soneson, 2014, Vieth et al., 2017, Benidt and Nettleton, 2015, Gerard, 2020, Zappia et al., 2017]. Some results were less expected from the original purpose of the methods. compcodeR and powsimR were developed mainly for differential-expression benchmarking and power analysis, yet neither gave the strongest pooled DE recovery here; compcodeR instead showed particularly strong repeated-run stability. SPsimSeq had the highest pooled recall but did not consistently perform best on the dependence metrics, despite being designed to retain reference-informed correlation structure. Overall, these results show that the purpose for which a simulator was developed does not fully determine how it performs across other properties of the simulated data.

Co-expression analyses further emphasized these differences. Before network topology could be evaluated, several simulated datasets failed to produce usable networks under the predefined construction criteria. compcodeR and powsimR generated near-zero-density networks across the evaluable reference datasets, and Splatter did so for three of the four evaluable datasets. Among simulators producing usable networks throughout, SimSeq most closely reproduced the reference network topology, although individual topology metrics favoured different simulators. Functional-coherence analyses similarly showed that plausible marginal expression patterns did not necessarily imply preservation of biologically meaningful network organization. Network evaluability and network fidelity should therefore be interpreted as separate aspects of simulation performance. At the same time, these findings remain conditional on the network construction strategy adopted here, since density, modularity, clustering, and related graph summaries depend on gene selection and edge-definition procedures [Langfelder and Horvath, 2008, Ballouz et al., 2015].

Repeated-run stability provided a complementary perspective. Unlike the other evaluation categories, Category D quantified reproducibility of simulator outputs rather than similarity to the reference data. compcodeR had the lowest pooled replicate-variation score, whereas seqgendiff, despite showing strong agreement for marginal and dependence fidelity, exhibited greater variability across repeated runs. This contrast demonstrates that reproducibility and biological fidelity represent different properties of simulated data. A simulator may generate highly reproducible outputs that differ systematically from the reference data, whereas another may exhibit greater stochastic variation while reproducing the desired biological characteristics more faithfully. Stability should therefore be interpreted as a complementary property of simulator behaviour rather than another measure of biological realism.

Differential-expression recovery further illustrates the importance of evaluating downstream analytical objectives directly. Although seqgendiff and SimSeq achieved the same pooled Category E score, the supporting evidence differed because seqgendiff was evaluable across all reference datasets whereas the SimSeq result reflected a single evaluable dataset. Individual recovery metrics also favoured different simulators, with SPsimSeq achieving the highest pooled recall and compcodeR the highest pooled Jaccard overlap. These findings indicate that successful preservation of marginal distributions or dependence structure alone cannot be assumed to guarantee recovery of downstream differential-expression signals.

Benchmark interpretation also depends on valid-output coverage. Simulation failure, undefined downstream statistics, and degenerate co-expression networks represent different outcomes and should not be interpreted equivalently. Except for the predefined network-usability criterion used in Category C, unavailable results were retained as missing values and reported separately from fidelity scores. This approach distinguishes biological performance from computational evaluability and avoids penalizing simulators for outcomes that cannot be meaningfully interpreted. Consequently, both performance and evaluability should be considered when comparing simulation methods.

Several limitations should be acknowledged. The five reference datasets provide biological and technical diversity but do not represent the full range of bulk RNA-seq study designs. All were human datasets, so performance in other organisms was not evaluated directly. More generally, simulator behaviour may depend on study characteristics such as sample size, sequencing depth, library size, expression variability, batch effects, and other experimental conditions. The primary benchmark used the 3000 most variable genes, although analyses with larger gene sets gave similar qualitative results for BRCA. Network results also depend on gene selection and graph-construction choices, and alternative definitions could change network evaluability or topology summaries. Finally, the simulator versions and parameter settings represent the configurations evaluated here rather than every possible implementation of each method.

The rank-normalized scoring framework also has limitations. Ranking places heterogeneous metrics on a common scale but does not retain the magnitude of performance differences. Likewise, the equal-weight composite combines five conceptually different evaluation targets and is therefore treated as a secondary descriptive summary, while category-specific results remain the main basis for interpretation. Dataset-bootstrap resampling with 2000 replicates also showed substantial overlap in the rank uncertainty of the two leading methods: seqgendiff had a median rank of 1 with a 2.5th–97.5th percentile interval of 1–2, while SimSeq had a median rank of 2 with the same interval. Full rank-uncertainty results are reported in Supplementary Section A8.

The broader contribution of this study extends beyond the seven simulators evaluated here. Rather than asking whether a simulator is generally realistic, the benchmark shows that simulator performance should be evaluated according to the statistical and biological properties required for the intended downstream analysis. Marginal count behaviour, dependence structure, co-expression topology, repeated-run stability, and differential-expression recovery provide complementary information, and preserving one does not necessarily imply preservation of another. Future bulk RNA-seq simulators may therefore benefit from modelling marginal distributions and dependence more explicitly while retaining sufficient joint structure for network and other multivariate analyses. More generally, simulation benchmarks should report which properties of the reference data are preserved, how reproducibly those properties are generated, and whether the resulting data remain usable for the downstream analyses they are intended to support. A multidimensional, task-specific framework therefore provides a more informative basis for simulator assessment and selection than a universal ranking.

## 5 Conclusion

This study evaluated seven bulk RNA-seq simulators across five reference datasets using five complementary evaluation dimensions. Simulator performance varied with both the property being evaluated and the reference dataset. seqgendiff performed best for marginal fidelity (Category A) and dependence structure (Category B), SimSeq for network topology among methods pro-ducing usable networks, and compcodeR for repeated-run stability. Differential-expression recovery produced a pooled tie between seqgendiff and SimSeq, although seqgendiff had substantially broader dataset coverage. These results show that no single summary adequately describes RNA-seq simulation performance. Agreement in marginal count behaviour did not ensure preservation of dependence, network structure, stability, or downstream differential-expression recovery. Simulator selection should therefore be guided by the properties required for the intended analysis, with performance, output coverage, and reproducibility considered together. The benchmark provides a task-specific framework for making these comparisons and identifies dependence and co-expression structure as important targets for further development of bulk RNA-seq simulation methods.

## A Benchmark Supporting Analyses

### A1 Dataset details and preprocessing

Five real bulk RNA-seq datasets were used as reference datasets: TEP (GSE68086), NSCLC (GSE239514), BRCA (TCGA-BRCA through recount3), AD (SRP181886 through recount3), and LUAD (TCGA-LUAD through recount3). The analyzed cohorts contained two biological groups and differed in sample size, sequencing depth, sparsity, and biological setting.

For BRCA, 100 normal and 100 tumour samples were selected without replacement using seed 42. For LUAD, 50 normal and 50 tumour samples were selected in the same manner using seed 42. For TEP, 50 control and 50 cancer samples were retained. The full processed AD cohort was used, consisting of 70 control and 80 AD samples. All 25 eligible NSCLC tumour-draining lymph-node samples were retained, comprising 13 CRT− and 12 CRT+ samples. In GSE239514, CRT+ denotes patients who received neoadjuvant chemoradiotherapy before radical surgery and lymph-node dissection, whereas CRT− denotes patients who underwent surgery without neoadjuvant chemoradiotherapy.

Source-specific expression filtering was performed before the common benchmark filter. BRCA retained genes with at least 10 counts in at least three samples; TEP retained genes with nonzero counts in at least three samples; AD retained genes with at least 10 counts in at least ten samples; and NSCLC retained genes with at least 10 counts in at least three samples. LUAD used the recount3-processed count matrix after gene-identifier harmonization. After these source-specific steps, the same reference filter was applied to every dataset: genes were required to have a total count greater than 20 and a mean count greater than 1. Counts exceeding 10^7^ were truncated, all-zero genes were removed, and gene identifiers were harmonized before variance-based selection.

For gene selection, the retained count matrices were TMM-normalized and transformed to log counts per million (logCPM) with a prior count of 1. Genes were ranked by row-wise logCPM variance. The top 3000 genes were retained for the primary count-level benchmark. The top 1000 genes under the same variance ranking were used for co-expression network analyses. Raw count values for the selected genes, rather than the transformed values used for ranking, were supplied to the simulators and used for count-level evaluation.

**Table A1:** Characteristics of the five reference datasets after preprocessing and selection of the 3000 most-variable genes. Median library size is the median total count across the retained genes, and zero fraction is the proportion of zero entries in the resulting count matrix.

| **Dataset** | **Biological setting** | **Samples** | **Group balance** | **Median library size** | **Zero fraction** |
|---|---|---|---|---|---|
| TEP | Platelet, cancer versus control | 100 | 50 control, 50 cancer | 1,974,174 | 0.244 |
| NSCLC | NSCLC tumour draining lymph nodes, neoadjuvant chemoradiotherapy status | 25 | 13 CRT-, 12 CRT+ | 10,084,357 | 0.039 |
| BRCA | Breast cancer | 200 | 100 normal, 100 tumor | 39,315,201 | 0.321 |
| AD | Alzheimer’s disease | 150 | 70 control, 80 AD | 38,550,377 | 0.392 |
| LUAD | Lung adenocarcinoma | 100 | 50 normal, 50 tumor | 39,137,814 | 0.137 |

**Table A2:**
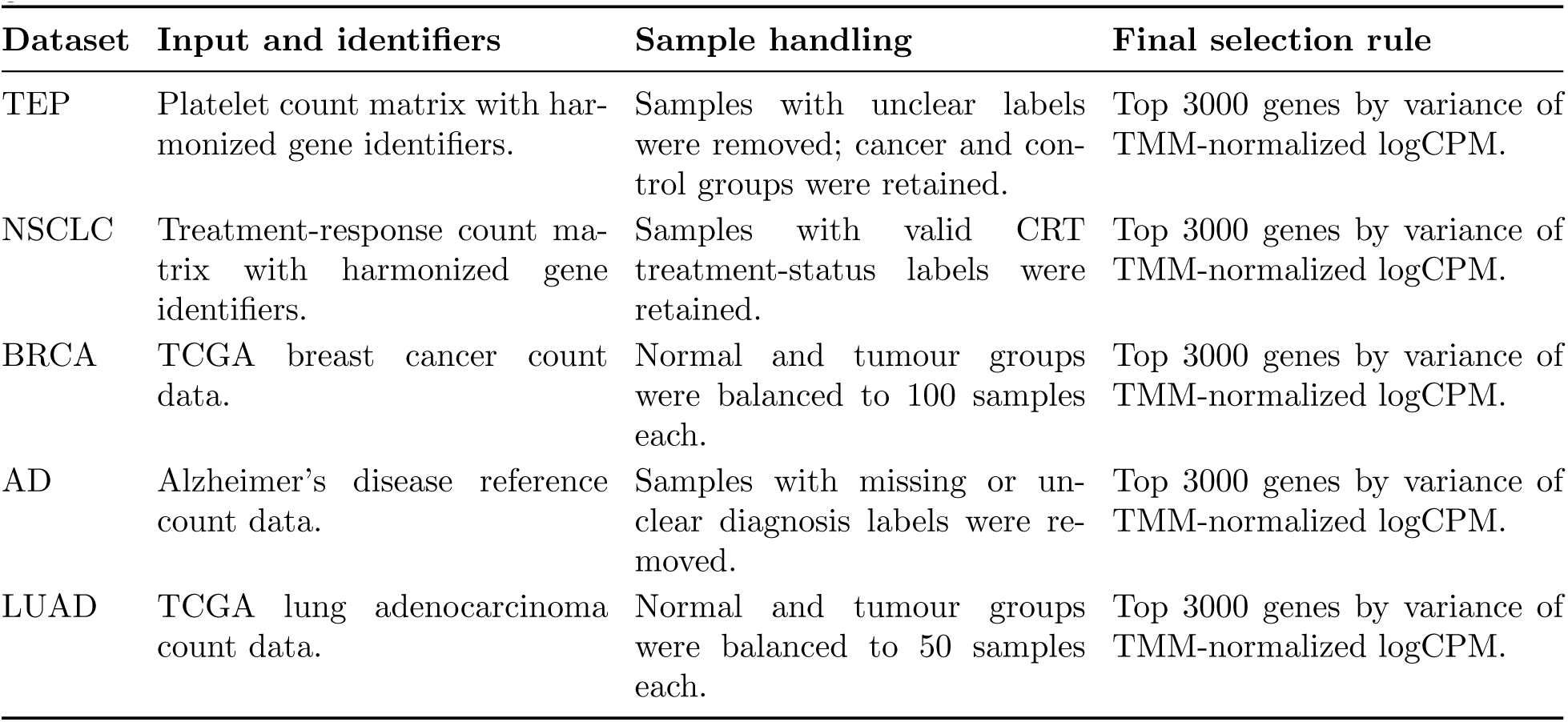
Dataset-specific preprocessing before the common variance-based selection of 3000 genes.

| **Dataset** | **Input and identifiers** | **Sample handling** | **Final selection rule** |
|---|---|---|---|
| TEP | Platelet count matrix with har-monized gene identifiers. | Samples with unclear labels were removed; cancer and con-trol groups were retained. | Top 3000 genes by variance of TMM-normalized logCPM. |
| NSCLC | Treatment-response count ma-trix with harmonized gene identifiers. | Samples with valid CRT treatment-status labels were retained. | Top 3000 genes by variance of TMM-normalized logCPM. |
| BRCA | TCGA breast cancer count data. | Normal and tumour groups were balanced to 100 samples each. | Top 3000 genes by variance of TMM-normalized logCPM. |
| AD | Alzheimer’s disease reference count data. | Samples with missing or un-clear diagnosis labels were re-moved. | Top 3000 genes by variance of TMM-normalized logCPM. |
| LUAD | TCGA lung adenocarcinoma count data. | Normal and tumour groups were balanced to 50 samples each. | Top 3000 genes by variance of TMM-normalized logCPM. |

Source-specific preprocessing was restricted to cleaning, label checking, identifier harmonization, group selection and the filtering needed to obtain an evaluable count matrix. The same variance-based selection procedure was then applied to all five datasets, providing a common 3000-gene dimension for the primary simulator comparison.

### A2 Metric definitions and scoring

The benchmark contains both primary scoring metrics and supporting diagnostics. Primary metrics contribute to the category scores, whereas supporting metrics are retained to describe additional properties of the simulated data without giving closely related or weakly discriminating quantities additional weight. Because the metrics have different units and numerical ranges, their raw values were not averaged directly. Simulators with valid results were ranked within each dataset and metric in the appropriate direction, and these ranks were mapped to a common scale from 0 to 1.

For a given dataset and scored metric, valid simulator outputs were ranked in ascending order when smaller values indicated stronger agreement and in descending order when larger values were preferred. Tied values received their average rank. If *r_m,k_* denotes the rank of simulator *m* on metric *k*, and *N_k_* is the number of simulators with valid values for that dataset–metric combination, the normalized score was

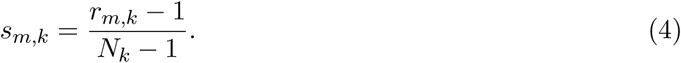

Thus, *s_m,k_*= 0 corresponds to the strongest valid result and *s_m,k_* = 1 to the weakest. Metrics with fewer than two valid simulator values were not used for comparative scoring.

Within Categories A, B, C and E, normalized scores were averaged across the primary metrics available for a simulator within each dataset. Final pooled scores for these four categories were obtained as the median of the corresponding dataset-level category scores across evaluable reference datasets. Missing simulator outputs remained unavailable rather than being assigned an artificial performance score.

Category C included an additional network-evaluability rule. AD was excluded from the primary Category C comparison because its reference network was not evaluable under the benchmark construction. For BRCA, LUAD, NSCLC and TEP, a simulated co-expression network with density ≤ 1 × 10^−4^ was classified as degenerate. A simulator that had been run successfully but produced such a network was assigned a Category C score of 1 for that dataset. This rule distinguishes failure to produce usable co-expression structure from a simulator output that was unavailable for other reasons.

Category D was derived separately from replicate-to-replicate variation in the available Category A, B, C and E measurements. For a replicate-based quantity *x*, variability was summarized using

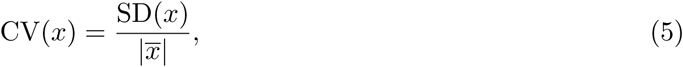

when the mean was non-zero and the quantity was defined. The available replicate-variation information was combined within each simulator–dataset combination and converted to a dataset-level normalized stability score. Lower values indicate less run-to-run variation. The final Category D score was the arithmetic mean of the five corrected dataset-level stability scores. Category D therefore measures reproducibility rather than agreement with a feature of the reference dataset.

For descriptive overall comparison, the five pooled category scores were given equal weight:

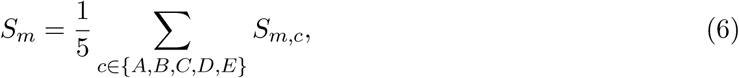

where *S_m,c_* is the pooled score for simulator *m* in category *c*. Because the categories represent different scientific targets, this composite was treated as a secondary summary rather than as the primary basis for interpreting simulator performance.

#### Worked scoring example

Table A3 illustrates the normalization using mean–variance ISE for the AD dataset. All seven simulators had valid values, so *N_k_*= 7.

**Table A3:**
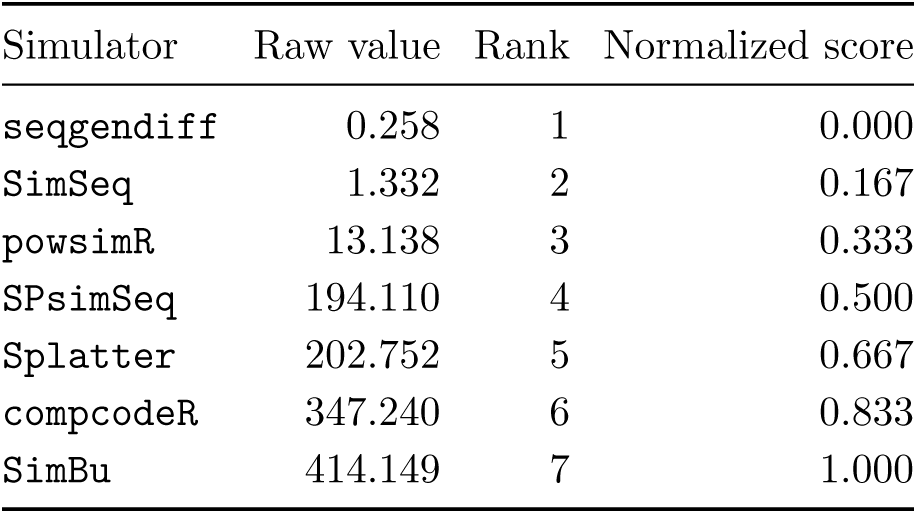
Worked example of rank normalization using mean–variance ISE in the AD dataset. Lower raw values indicate closer agreement with the reference.

With seven valid simulator values, the denominator is 7 − 1 = 6. Rank 1 therefore maps to 0*/*6 = 0, rank 2 to 1*/*6 = 0.167, and rank 7 to 6*/*6 = 1. When fewer simulators have valid values, *N_k_* changes accordingly rather than retaining a fixed seven-simulator denominator.

### A3 Full raw metric values

The complete raw metric summaries for Categories A, B, C and E are provided as a machine-readable file. Each row identifies the simulator, reference dataset, metric and evaluation category together with the mean value, standard deviation and number of contributing simulation replicates. These values provide the direct link between replicate-level benchmark measurements and the category summaries reported in the main text. The number of contributing replicates is retained because some simulator–dataset–metric combinations were not evaluable in every run. Reporting this quantity separately distinguishes summaries based on complete replicate sets from those based on fewer valid outputs without assigning artificial values to unavailable runs.

Category D is not represented as an additional raw fidelity metric in this file because it is derived from replicate-to-replicate variation in the Category A, B, C and E measurements. The stability inputs and corrected dataset-level Category D scores are provided separately with the deposited stability outputs. The deposited raw-metric summary is

FINAL_raw_metrics_mean_sd.csv.

**Table A4:**
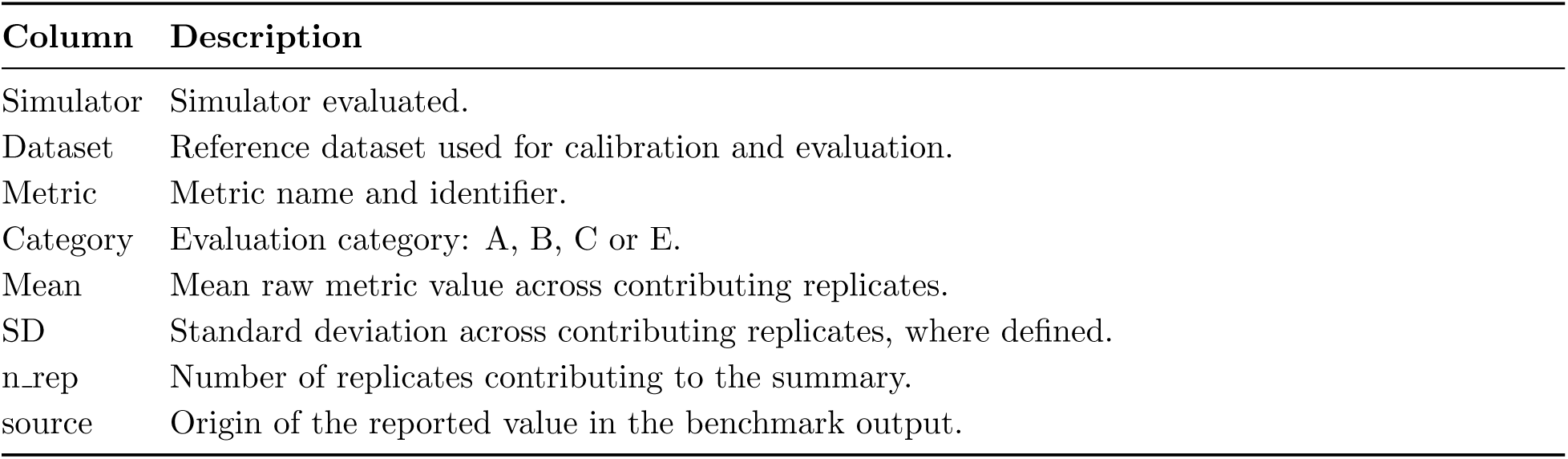
Column structure of the machine-readable raw-metric summary.

| **Column** | **Description** |
|---|---|
| Simulator | Simulator evaluated. |
| Dataset | Reference dataset used for calibration and evaluation. |
| Metric | Metric name and identifier. |
| Category | Evaluation category: A, B, C or E. |
| Mean | Mean raw metric value across contributing replicates. |
| SD | Standard deviation across contributing replicates, where defined. |
| n_rep | Number of replicates contributing to the summary. |
| source | Origin of the reported value in the benchmark output. |

### A4 Metric-wise rankings

Metric-wise summaries retain information that can be obscured by a category-level score. They are reported for Categories A, B, C and E and include both the primary metrics used in scoring and the supporting diagnostics defined in Supplementary Section A2. Category D is not included in these metric-wise fidelity rankings because it is derived separately from replicate-to-replicate variation.

The metric-level results show that methods need not perform similarly across different measurements of the same broad data property. This was most apparent for dependence. Gene– gene Frobenius distance, sample–sample Frobenius distance, Procrustes *M* ^2^, PVE correlation and the supporting RV coefficient produced different simulator orderings. The pooled category score therefore summarizes several related measurements rather than a single definition of dependence preservation.

The complete metric-wise results are in the associated file:

FINAL_permetric_pooled_metricwise.csv

Category-specific files are provided for Categories A, B, C and E. Supporting metrics remain in these files for interpretation but do not contribute additional weight to the category scores.

**Table A5:** Column structure of the machine-readable metric-wise ranking summary. Category D is reported separately through the stability analysis.

| **Column** | **Description** |
|---|---|
| Category | Evaluation category: A, B, C or E. |
| Metric | Metric name and identifier. |
| rank | Simulator rank for the pooled metric summary. |
| Simulator | Simulator evaluated. |
| Value | Pooled metric summary across datasets with valid values. |
| n_datasets_defined | Number of reference datasets contributing to the pooled value. |
| Lower_better | Indicator for whether smaller raw values correspond to stronger agreement. |

### A5 Per-dataset category rankings

Dataset-level category scores retain variation that is hidden by the pooled summaries. Lower values indicate stronger performance within a given dataset and category. Because the scores are rank-normalized within a reference setting, a score of 0 denotes the strongest observed result in that comparison and should not be interpreted as zero numerical discrepancy from the reference.

The final dataset-level results incorporate two corrections used throughout the main analysis. AD is excluded from Category C because its reference network was not evaluable under the benchmark construction, and Category D uses the corrected replicate-stability scores. The leading simulator therefore changes across both datasets and evaluation targets (Table A6).

**Table A6:** Lowest dataset-level category scores after the final Category C and Category D corrections. Values in parentheses are rank-normalized category scores. Ties are shown explicitly. AD was not evaluated for Category C.

| **Dataset** | **A: Marginal** | **B: Dependence** | **C: Network** | **D: Stability** | **E: DE recovery** |
|---|---|---|---|---|---|
| TEP | seqgendiff (0.000) | seqgendiff / Splatter (0.292) | SimSeq (0.333) | compcodeR (0.000) | seqgendiff (0.000) |
| NSCLC | SimSeq (0.167) | seqgendiff / Splatter (0.333) | SimSeq (0.000) | powsimR (0.000) | seqgendiff / SimSeq (0.167) |
| BRCA | seqgendiff (0.000) | compcodeR / seqgendiff (0.292) | SimSeq (0.297) | compcodeR (0.000) | SPsimSeq (0.000) |
| AD | seqgendiff (0.067) | SimSeq (0.208) | Not evaluated | compcodeR (0.000) | seqgendiff (0.000) |
| LUAD | seqgendiff (0.040) | seqgendiff (0.350) | seqgendiff (0.000) | Splatter (0.000) | SPsimSeq (0.000) |

The dataset-level pattern explains why the pooled results should not be read as fixed simulator orderings. For example, seqgendiff had the lowest marginal score in four datasets but not NSCLC. Dependence contained several dataset-specific ties, whereas the network category favoured SimSeq in three of the four evaluable datasets and seqgendiff in LUAD. Stability showed a different pattern again: compcodeR ranked first in TEP, BRCA and AD, powsimR in NSCLC, and Splatter in LUAD. The associated corrected dataset-level score file is named FINAL category scores per dataset FINAL.csv. These are the same values used to generate the final dataset-specific ranking figure and the dataset-bootstrap analysis.

### A6 Valid-output availability and metric coverage

Availability was recorded separately from fidelity because an unavailable simulation, an undefined downstream statistic and a valid but poorly matching result represent different outcomes. Coverage therefore describes whether an expected result could be evaluated; it is not itself a measure of similarity to the reference data.

For Categories A, B and E, unavailable simulator–dataset combinations were left missing rather than assigned a poor score. Category C required additional handling because a simulator could generate a valid count matrix but still produce a co-expression network too sparse for meaningful topology summaries. In that case, the failure to produce usable network structure was treated as part of Category C performance rather than as ordinary missingness.

**Table A7:** Column structure of the machine-readable coverage summary. Category D is handled separately through the corrected replicate-stability analysis.

| **Column** | **Description** |
|---|---|
| Category | Evaluation category: A, B, C or E. |
| Simulator | Simulator evaluated. |
| Coverage | Proportion of expected evaluable dataset–metric combinations with valid output. |

Several specific availability rules affect interpretation of the benchmark (Table A8). Structural exclusions, such as AD in Category C, are outside the corresponding coverage denominator rather than being counted as simulator failures.

For Category C, a simulated network with density *≤* 1 *×* 10*^−^*^4^ was classified as degenerate. Such a result indicates that the simulator produced an expression matrix but that the resulting co-expression structure could not support the predefined topology comparison. These cases were therefore assigned the worst Category C score of 1 rather than removed from the analysis.

This distinction prevents two different situations from being conflated. A missing simulator output provides no evidence about fidelity for that dataset, whereas a generated count matrix that produces a degenerate network provides direct evidence that the simulated data did not support the network analysis under the benchmark rule. Coverage and performance are therefore reported together but interpreted separately. The deposited coverage and network-validity files record the corresponding availability indicators and Category C usability decisions.

**Table A8:**
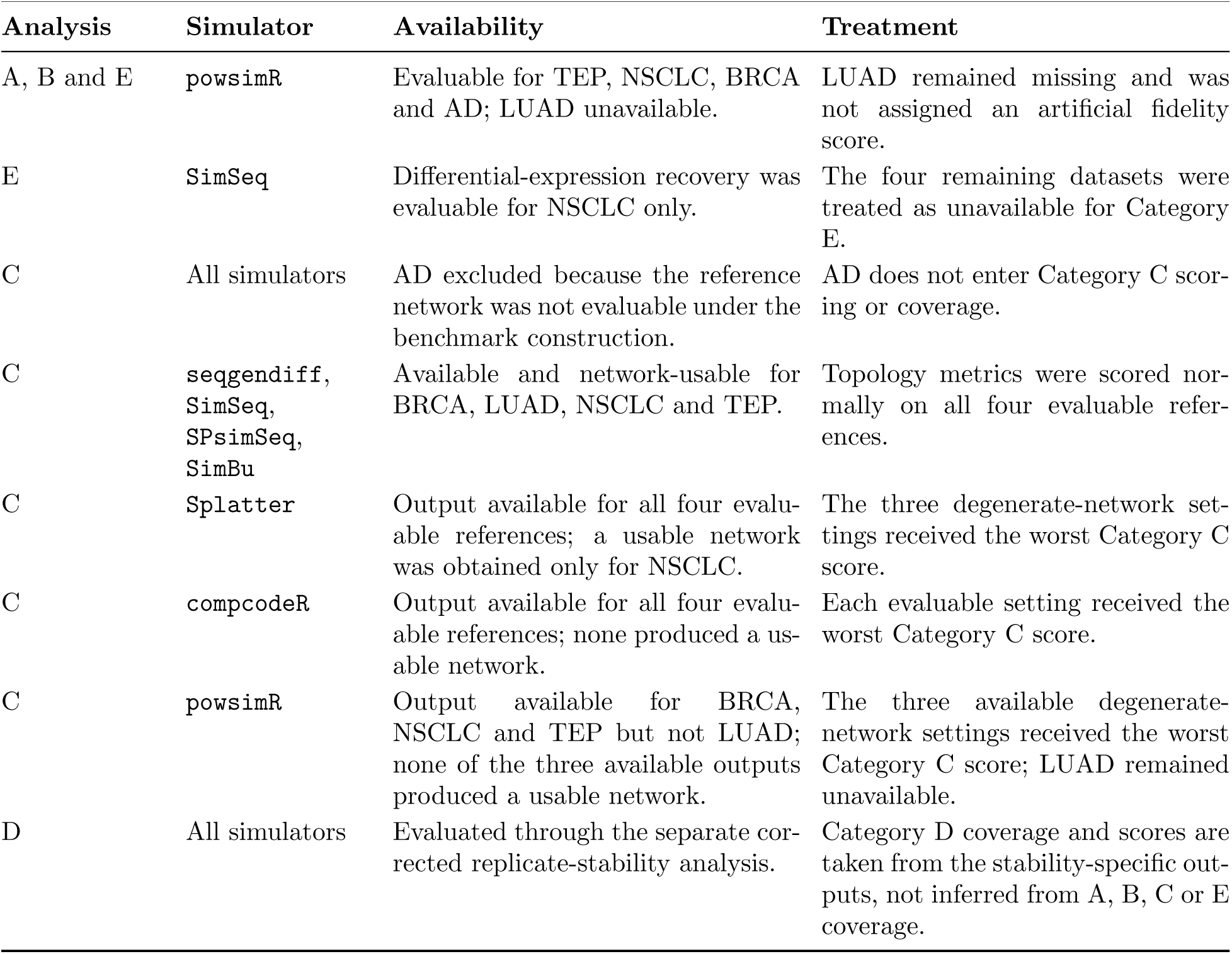
Principal output-availability and network-usability rules in the final benchmark. “Available” indicates that simulator output was present for the analysis; “usable” in Category C additionally requires a non-degenerate co-expression network.

| Analysis | Simulator | Availability | Treatment |
|---|---|---|---|
| A, B and E | powsimR | Evaluable for TEP, NSCLC, BRCA and AD; LUAD unavailable. | LUAD remained missing and was not assigned an artificial fidelity score. |
| E | SimSeq | Differential-expression recovery was evaluable for NSCLC only. | The four remaining datasets were treated as unavailable for Category E. |
| C | All simulators | AD excluded because the reference network was not evaluable under the benchmark construction. | AD does not enter Category C scor-ing or coverage. |
| C | seqgendiff, SimSeq, SPsimSeq, SimBu | Available and network-usable for BRCA, LUAD, NSCLC and TEP. | Topology metrics were scored nor-mally on all four evaluable refer-ences. |
| C | Splatter | Output available for all four evalu-able references; a usable network was obtained only for NSCLC. | The three degenerate-network set-tings received the worst Category C score. |
| C | compcodeR | Output available for all four evalu-able references; none produced a us-able network. | Each evaluable setting received the worst Category C score. |
| C | powsimR | Output available for BRCA, NSCLC and TEP but not LUAD; none of the three available outputs produced a usable network. | The three available degenerate-network settings received the worst Category C score; LUAD remained unavailable. |
| D | All simulators | Evaluated through the separate cor-rected replicate-stability analysis. | Category D coverage and scores are taken from the stability-specific out-puts, not inferred from A, B, C or E coverage. |

### A7 Pairwise dissociability

Pairwise dissociability was used to assess whether the evaluation categories produced similar simulator orderings. Within each reference dataset, Kendall’s *τ* was calculated between the simulator orderings induced by each pair of Categories A, B, C, and E, using only simulators with valid scores in both categories. Category D was excluded because it measures replicate-to-replicate stability rather than agreement with a property of the reference data. Values of *τ* closer to 1 indicate more similar simulator orderings, whereas lower values indicate greater disagreement between evaluation targets. The dataset-specific correlations were summarized by their mean and median. Because Category C was not evaluable for AD, all comparisons involving Category C were based on BRCA, LUAD, NSCLC, and TEP only.

The highest agreement was observed between marginal fidelity and dependence (A–B), with a mean *τ* of 0.585 and a median of 0.552 across the five reference datasets. Marginal fidelity and DE recovery (A–E) showed moderate agreement (mean 0.483; median 0.500), while dependence and DE recovery (B–E) showed lower agreement (mean 0.386; median 0.357). Comparisons involving network topology were generally less concordant. Marginal fidelity and network topology (A–C) had a mean *τ* of 0.389 and a median of 0.327, whereas network topology and DE recovery (C–E) had a mean of 0.352 and a median of 0.365 across the four evaluable datasets. The lowest agreement occurred between dependence and network topology (B–C), with a mean *τ* of 0.129 and a median of 0.080. Thus, simulators that ranked similarly for dependence preservation did not necessarily rank similarly for co-expression network topology.

**Table A9:**
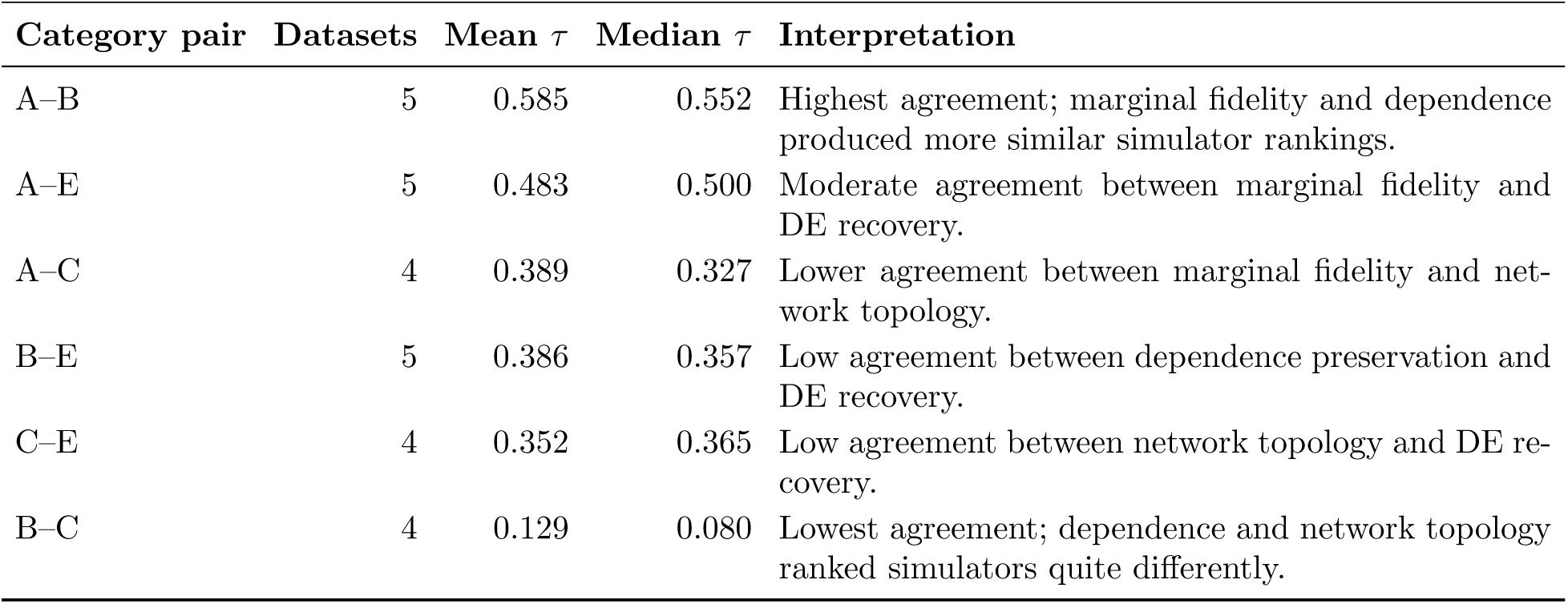
Pairwise agreement between simulator rankings across evaluation categories, summarized across reference datasets. Lower Kendall’s *τ* indicates greater dissociability between evaluation targets. Category D is excluded because it measures replicate stability rather than agreement with the reference data. Comparisons involving Category C exclude AD because the reference network was not evaluable.

| **Category pair** | **Datasets** | **Mean τ** | **Median τ** | **Interpretation** |
|---|---|---|---|---|
| A–B | 5 | 0.585 | 0.552 | Highest agreement; marginal fidelity and dependence produced more similar simulator rankings. |
| A–E | 5 | 0.483 | 0.500 | Moderate agreement between marginal fidelity and DE recovery. |
| A–C | 4 | 0.389 | 0.327 | Lower agreement between marginal fidelity and net-work topology. |
| B–E | 5 | 0.386 | 0.357 | Low agreement between dependence preservation and DE recovery. |
| C–E | 4 | 0.352 | 0.365 | Low agreement between network topology and DE re-covery. |
| B–C | 4 | 0.129 | 0.080 | Lowest agreement; dependence and network topology ranked simulators quite differently. |

These correlations are interpreted descriptively. Only five reference datasets were available, reduced to four for comparisons involving Category C, and the number of simulators contributing to an individual correlation could be smaller when a category score was unavailable. The analysis therefore supports the view that the evaluation categories contain partly distinct ranking information, but the individual Kendall correlations are not treated as population-level estimates or formal evidence of category independence.

The corrected dataset-level dissociability results and their across-dataset summaries are in the associated files:

FINAL_dissociability_allpairs_FINAL.csv

FINAL_dissociability_summary_FINAL.csv

### A8 Dataset-bootstrap rank uncertainty

For each simulator, the bootstrap rank distribution was summarized by its median rank together with the 2.5th and 97.5th percentile ranks. These intervals describe the sensitivity of the aggregate ordering to reference dataset composition rather than confidence intervals for a population parameter.

**Table A10:** Overall ranking with dataset-bootstrap rank intervals. The overall score is the equal-weight composite of the five pooled category scores, with lower values indicating stronger aggregate performance. Bootstrap intervals summarize sensitivity to reference-dataset composition.

| **Simulator** | **Overall score** | **Median rank** | **Lower rank** | **Upper rank** | **Valid bootstraps** |
|---|---|---|---|---|---|
| seqgendiff | 0.351 | 1 | 1 | 2 | 2000 |
| SimSeq | 0.356 | 2 | 1 | 2 | 1358 |
| SPsimSeq | 0.512 | 3 | 2 | 4 | 2000 |
| compcodeR | 0.562 | 4 | 2 | 7 | 2000 |
| powsimR | 0.632 | 5 | 3 | 7 | 1976 |
| Splatter | 0.663 | 5 | 3 | 7 | 2000 |
| SimBu | 0.733 | 7 | 5 | 7 | 2000 |

The bootstrap results showed substantial overlap between the two leading simulators. seqgendiff had a median rank of 1 with a 2.5th–97.5th percentile interval of 1–2, while SimSeq had a median rank of 2 with the same interval. SPsimSeq had a median rank of 3 with an interval of 2–4. Greater variation in aggregate rank was observed for the middle-ranked simulators, with compcodeR, powsimR, and Splatter showing broad overlapping intervals. SimBu remained toward the lower end of the aggregate ordering, with a median rank of 7 and an interval of 5–7.

The number of valid bootstrap iterations is also reported because not every simulator was evaluable in every bootstrap sample. SimSeq contributed to 1358 of the 2000 bootstrap replicates because its Category E result was available only for NSCLC. powsimR contributed to 1976 valid replicates, reflecting its incomplete LUAD coverage.

### A9 Pairwise statistical comparisons

We used pairwise statistical tests to examine differences between individual simulators. These comparisons provided supporting evidence and were not used to calculate category scores or determine the overall rankings. Within each reference dataset and metric, we compared pairs of simulators using the Wilcoxon rank-sum test (Mann–Whitney U) on their ten replicate values. Cliff’s delta measured the direction and size of the differences between their metric distributions. Positive values indicate that the first simulator tended to have larger metric values, while negative values indicate the opposite. Whether larger values represent better performance depends on the metric.

We applied the Benjamini–Hochberg correction once across all within-dataset pairwise tests, including every dataset, metric, and simulator pair. The adjusted *p*-values therefore account for multiple testing across the full family of these comparisons. We also used paired Wilcoxon signed-rank tests to compare simulator metric summaries across reference datasets. These tests examined whether the differences between two simulators were consistent across the available datasets. Only datasets with valid results for both simulators were included. Category C excluded AD, and unavailable simulator results, including powsimR on LUAD, were not assigned artificial values.

The across-dataset tests included at most five paired reference datasets, giving them limited statistical power. Most comparisons did not show statistically significant differences after multiple-testing adjustment. This does not establish equivalent simulator performance. We therefore based the main benchmark findings on the observed metric values, category scores, and consistency across datasets rather than on statistical significance alone. The supporting files are:

pairwise_tests.csv

pairwise_signedrank_across_datasets.csv

**Table A11:** Column structure of the deposited pairwise-comparison outputs. Not every column occurs in both files because within-setting and across-dataset comparisons answer different questions.

| Column | Description |
|---|---|
| Dataset | Reference dataset for comparisons performed within a reference setting. |
| Metric | Benchmark metric being compared. |
| Sim1, Sim2 | Pair of simulators compared. |
| p_value | Nominal pairwise *p*-value, where applicable. |
| p_adj_BH | Benjamini–Hochberg adjusted *p*-value. |
| cliffs_delta | Cliff’s delta effect-size estimate, where available. |
| n_datasets | Number of paired reference datasets contributing to an across-dataset comparison. |
| signedrank_p | Wilcoxon signed-rank *p*-value across paired datasets. |

### A10 Composite ranking and sensitivity analyses

The primary interpretation of the benchmark is based on the five evaluation categories rather than on a single overall score. Nevertheless, a composite summary is useful for describing broad patterns across simulators when its limitations are recognised explicitly. The composite ranking was therefore constructed as a descriptive summary rather than as a universal measure of simulator quality. The pooled category scores used in the composite analysis are reproduced in Table A12. Categories A, B, C and E were pooled across reference datasets using the median of the dataset-level category scores, whereas the corrected stability category (Category D) was pooled using the arithmetic mean of the corrected dataset-level stability scores. All five categories contributed equal weight to the composite score.

The equal-weight composite placed seqgendiff first at 0.351 and SimSeq second at 0.356, a difference of only 0.005. Their category profiles were nevertheless different: seqgendiff had lower marginal and dependence scores, whereas SimSeq had the lower network-topology score. Similar composite values therefore do not imply similar simulator behaviour. The dataset-bootstrap analysis (Supplementary Section A8) supports this interpretation that the aggregate ordering is sensitive to reference-dataset composition. The two leading simulators had overlapping rank intervals, and the interval for SPsimSeq extended to rank 2. Broader intervals were observed among several middle-ranked methods, reflecting changes in aggregate ordering across resampled combinations of the five reference datasets.

**Table A12:** Final pooled category scores and equal-weight composite scores. Lower values indicate stronger aggregate performance under the benchmark scoring framework.

| **Rank** | **Simulator** | **A** | **B** | **C** | **D** | **E** | **Composite** |
|---|---|---|---|---|---|---|---|
| 1 | seqgendiff | 0.040 | 0.292 | 0.458 | 0.800 | 0.167 | 0.351 |
| 2 | SimSeq | 0.200 | 0.417 | 0.297 | 0.700 | 0.167 | 0.356 |
| 3 | SPsimSeq | 0.567 | 0.550 | 0.594 | 0.600 | 0.250 | 0.512 |
| 4 | compcodeR | 0.720 | 0.500 | 1.000 | 0.133 | 0.458 | 0.562 |
| 5 | powsimR | 0.567 | 0.458 | 1.000 | 0.533 | 0.600 | 0.632 |
| 6 | Splatter | 0.600 | 0.450 | 1.000 | 0.367 | 0.900 | 0.663 |
| 7 | SimBu | 0.867 | 0.850 | 0.683 | 0.367 | 0.900 | 0.733 |

Several aspects of the benchmark also affect interpretation of the composite score. Category C excluded AD because the reference network was not evaluable under the benchmark construction, and Category D measures replicate reproducibility rather than similarity to a reference dataset. Likewise, coverage differed among simulators, with SimSeq contributing differential-expression results only for NSCLC and powsimR remaining unavailable for LUAD. These differences are reflected in the category-specific summaries but cannot be represented fully by a single aggregate score.

For these reasons, the composite ranking is presented as a descriptive overview that complements, rather than replaces, the task-specific benchmark. The principal conclusions of the study are therefore based on the individual evaluation categories, dataset-specific behaviour and output coverage.

### A11 Software versions, simulator classes, and settings

#### A11.1 Simulator classes and modeling assumptions

The seven simulators benchmarked in this study differ substantially in their statistical assumptions and data-generation strategies. These differences provide useful context for interpreting the benchmark results but should not be regarded as a complete explanation of performance. A first distinction concerns how information from the reference dataset is used. SimSeq and seqgendiff generate new observations by resampling or transforming empirical expression profiles, thereby preserving many characteristics of the observed data without specifying a fully parametric model. In contrast, compcodeR, powsimR, Splatter, and SPsimSeq estimate statistical models from the reference data and generate new counts from the fitted distributions. SimBu combines simulated cell-type profiles to produce bulk-like expression data.

The benchmark results are broadly consistent with these modeling strategies. Methods that retained stronger empirical structure generally performed well for marginal distributions and dependence, whereas model-based approaches showed greater variation across evaluation categories. However, no modeling strategy was uniformly superior. For example, compcodeR produced the lowest corrected stability score but ranked less favorably for network topology, while SimSeq showed the strongest network preservation among simulators producing usable co-expression networks despite not having the lowest marginal or dependence scores.

All simulators were evaluated using their recommended workflows and parameter settings for bulk RNA-seq data whenever possible. No simulator was tuned to optimize performance for a particular benchmark category. Consequently, the reported results represent comparisons between representative implementations rather than theoretical limits of the underlying statistical models.

**Table A13:** Simulator settings and random-number handling used in the main benchmark. Only selected non-default settings are shown. Here, *di* is the dataset index and *r* is the replicate index.

| Simulator | Seed strategy | Key parameters | Notes |
|---|---|---|---|
| compcodeR | set.seed(123) | minfact=0.7, maxfact=1.4<br>effect.size=1.5<br>fraction.upregulated=1 | Fixed configuration. |
| powsimR | set.seed(7000+10*di+r)<br>setup.seed=5000 | GeneFilter=0.05<br>SampleFilter=5<br>Distribution="NB"<br>Normalisation="MR" | Global seed varies by dataset and replicate; setup seed is fixed. |
| SimSeq | No explicit seed in wrapper. | sort.method="unpaired"<br>k.ind=max(3, min(n1,n2, round(0.3*min(n1,n2))))<br>n.genes=G<br>n.diff=round(0.05*G) | Binary group labels (0/1). Normalization factors computed with edgeR and rescaled to mean 1. |
| seqgendiff | 3000+r (NSCLC)<br>2000+r (TEP) | rate=0.5<br>es=1.5<br>group_prop=0.5 | Dataset-specific seeds. |
| SPsimSeq | No explicit seed in wrapper. | genewiseCor=TRUE<br>lfc.thrld=0.5<br>group.config=c(0.5,0.5) | Reference-informed simulation. |
| Splatter | set.seed(8000+r) | dropout.type="none"<br>method="groups"<br>group.prob=c(0.5,0.5) | Replicate-specific seed. |
| SimBu | set.seed(600+r) | scenario="random"<br>scaling_factor="NONE"<br>ncells=100 | Replicate-specific seed. |

All analyses were performed using R version 4.4.0. Table A14 lists the principal software packages used for simulator fitting, benchmarking, downstream analyses, and figure generation.

**Table A14:** Software versions used in the benchmarking pipeline. Analyses were performed in R version 4.4.0 (2024-04-24). Simulator versions correspond to those used during benchmark generation. SimBu simulations were performed previously using the archived benchmark outputs and the package was not installed in the final computational environment used to regenerate summary tables and figures.

| **Software / package** | **Version** |
|---|---|
| R | 4.4.0 |
| edgeR | 4.4.2 |
| DESeq2 | 1.46.0 |
| limma | 3.62.2 |
| WGCNA | 1.74 |
| EGAD | 1.34.0 |
| SPsimSeq | 1.16.0 |
| Splatter | 1.30.0 |
| powsimR | 1.2.5 |
| SimSeq | 1.4.0 |
| compcodeR | 1.42.0 |
| seqgendiff | 1.2.4 |
| SimBu | Archived benchmark outputs |
| msigdbr | 26.1.0 |
| igraph | 2.3.1 |

### A12 Functional coherence of simulated co-expression networks

Functional coherence was evaluated as a supporting analysis using EGAD neighbour voting on ranked co-expression networks with MSigDB Hallmark gene sets. The analysis was applied to all seven benchmark simulators across the five reference datasets. For each real and simulated network, neighbour voting was used to predict gene-set membership from network neighbours, and AUROC was averaged across the evaluable Hallmark gene sets. An AUROC of 0.5 corresponds to chance-level discrimination. The number of evaluable gene sets varied across datasets, from 3 in AD to 42 in TEP. This analysis used a ranked co-expression network rather than the density-thresholded network used for Category C. It could therefore remain evaluable for simulator–dataset combinations for which the Category C topology metrics were undefined or degenerate. GBA results were treated as supporting evidence and were not included in the Category C score.

Across the five datasets, the real networks had a mean AUROC of 0.682. seqgendiff had a similar mean AUROC of 0.681 and the smallest mean absolute difference from the corresponding real networks (0.012). SimBu and SimSeq had mean AUROCs of 0.653 and 0.625, with mean absolute differences of 0.047 and 0.062, respectively. The remaining simulators had larger mean differences from the corresponding real-network values (Table A15). These results provide an additional description of functional structure in the simulated networks but were not used in simulator ranking.

**Table A15:** Functional coherence of real and simulated co-expression networks using EGAD neighbour voting. AUROC summarizes prediction of MSigDB Hallmark gene-set labels from network neighbours. Mean absolute gap is the average absolute difference from the corresponding real-network AUROC across the five reference datasets. The analysis was used as a supporting biological check and was not included in Category C scoring.

| **Source** | **Mean AUROC** | **Mean abs. gap from real** | **Datasets** |
|---|---|---|---|
| Real networks | 0.682 | – | 5 |
| seqgendiff | 0.681 | 0.012 | 5 |
| SimBu | 0.653 | 0.047 | 5 |
| SimSeq | 0.625 | 0.062 | 5 |
| compcodeR | 0.519 | 0.163 | 5 |
| powsimR | 0.505 | 0.177 | 5 |
| SPsimSeq | 0.493 | 0.188 | 5 |
| Splatter | 0.493 | 0.189 | 5 |

### A13 Additional experimental-design analyses

The main benchmark focused on how closely each simulator reproduced observed features of real bulk RNA-seq data. Two additional questions are relevant when simulations are used for study planning: how differential-expression detection power changes with sample size, and how sensitive the comparison is to the number of genes retained. These analyses are reported as supporting results rather than as additional benchmark categories.

The primary benchmark was performed using the top 3000 most-variable genes from each reference dataset. Additional analyses examined whether the major benchmark conclusions were sensitive to the number of simulated samples or the number of retained genes.

#### A13.1 Sample-size effects and detection power

The sample-size analysis was included as a supporting study-design experiment rather than as an exhaustive comparison of all seven simulators. Under a common external differential-expression workflow, compcodeR and Splatter were evaluated at increasing sample sizes using the BRCA and TEP reference settings. powsimR was evaluated separately through its package-specific power routine because power estimation is one of its intended uses. SimSeq, seqgendiff, SPsimSeq, and SimBu were not included in this additional experiment; their omission reflects the scope of this supporting analysis rather than a general limitation on their ability to generate data at larger sample sizes.

For compcodeR and Splatter, data were generated at 10, 15, 25, 50, and 100 samples per group using the BRCA and TEP reference settings, with five repeats at each size. Detection power was defined as the fraction of truly differential genes recovered at an adjusted *p*-value below 0.05. Power was evaluated with DESeq2, edgeR, and limma-voom so that the observed pattern was not tied to a single differential-expression method. Because the powsimR procedure uses different internal assumptions, its estimates are reported separately and are not treated as directly comparable with the shared external workflow.

Power increased with sample size for all three simulators, but the level differed across methods. compcodeR reached relatively high power and crossed the common 0.80 level at roughly 25 to 50 samples per group, depending on the dataset. Splatter increased more gradually and remained below 0.80 even at 100 samples per group, consistent with weaker differential-expression signal in this bulk RNA-seq setting. At 25 samples per group with DESeq2 on BRCA, detection power was 0.82 for compcodeR and 0.29 for Splatter; on TEP, the same setting gave 0.74 and 0.10. Through its own routine, powsimR showed power increasing from about 0.25 to 0.70 on BRCA and from 0.03 to 0.81 on TEP across 10 to 100 samples per group, crossing 0.80 only on TEP at the largest sample size.

**Table A16:** Detection power on BRCA at selected sample sizes per group. compcodeR and Splatter use the shared DESeq2 evaluation, whereas powsimR uses its package-specific power routine and is therefore shown as a supporting comparison. The complete sample-size trajectories, including the 15-sample setting, are shown in Figure A7.

| **Selected Samples per group** | **compcodeR** | **Splatter** | **powsimR (own routine)** |
|---|---|---|---|
| 10 | 0.66 | 0.14 | 0.25 |
| 25 | 0.82 | 0.29 | 0.47 |
| 50 | 0.91 | 0.44 | 0.58 |
| 100 | 0.97 | 0.53 | 0.70 |

Because these power estimates depend on how each simulator introduces differential signal, part of the difference reflects simulator-specific DE-generation behavior rather than sample size alone. The comparison is therefore interpreted as evidence that simulator choice can affect the power–sample-size relationship, not as an absolute power estimate for any one biological study. In practical terms, a power calculation based on one simulator may suggest a different sample size than a calculation based on another simulator.

#### A13.2 Gene-set-size sensitivity

The main benchmark used a fixed 3000-gene setting to keep the comparison dimension consistent across datasets and simulators. To check whether this choice influenced the comparative results, the marginal and dependence metrics were recomputed on BRCA using 5000, 10000, and 15000 genes for simulators that ran across that range. Absolute metric values changed with gene count, as expected, because larger gene sets include additional genes and can change distance-based summaries. However, the relative ordering of simulators was stable across the evaluated gene-set sizes: seqgendiff remained closest to the real structure, and the lower-ranked methods remained lower-ranked. These results indicate that the 3000-gene setting captured the same comparative pattern observed at larger gene-set sizes. Robustness across additional datasets and still-larger gene sets is left for future work.

## A14 Detailed category result tables

This section provides the detailed category-level tables supporting the summarized results in the main text. For each category, we report the pooled simulator ranking, metric-level leaders where applicable, and per-dataset leading simulators. These tables were moved to the supplementary material to keep the main text focused on task-specific interpretation.

### Category A: Marginal distribution fidelity

**Table A17:**
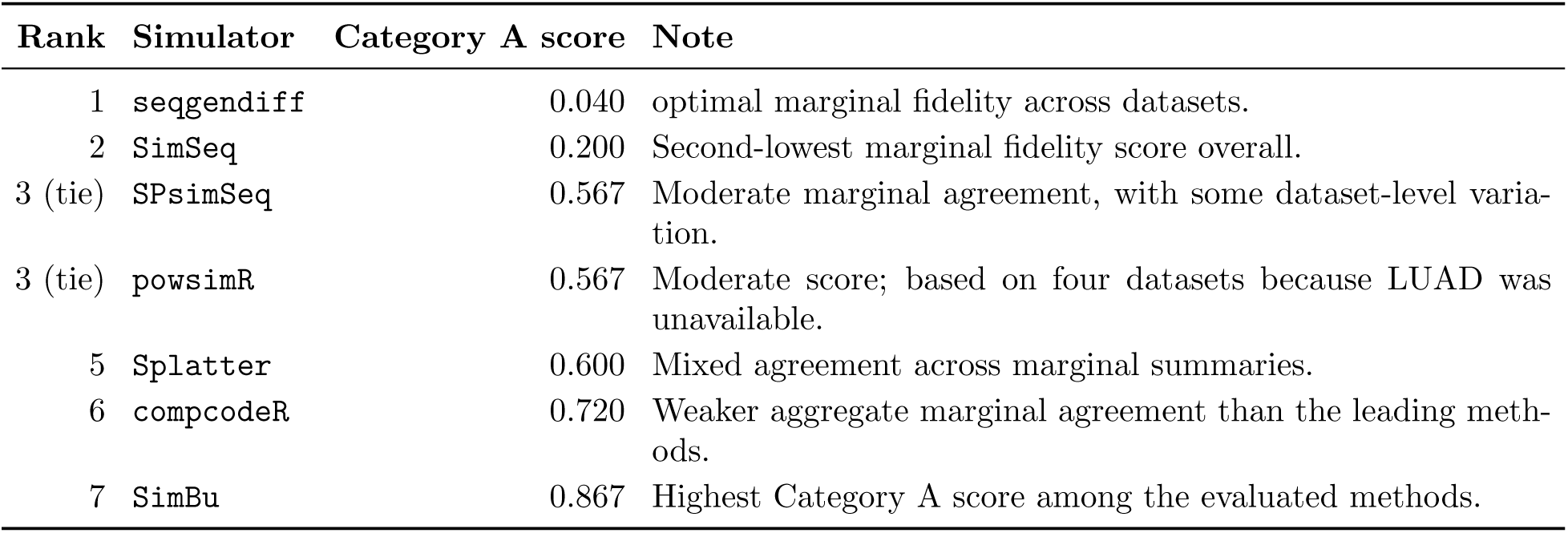
Category A ranking for marginal distribution fidelity. Scores are rank-normalized category summaries pooled across the benchmark datasets; lower values indicate stronger agreement. `powsimR’s` score is pooled over four of five datasets because no LUAD result was available.

| **Rank** | **Simulator** | **Category A score** | **Note** |
|---|---|---|---|
| 1 | seqgendiff | 0.040 | optimal marginal fidelity across datasets. |
| 2 | SimSeq | 0.200 | Second-lowest marginal fidelity score overall. |
| 3 (tie) | SPsimSeq | 0.567 | Moderate marginal agreement, with some dataset-level varia-tion. |
| 3 (tie) | powsimR | 0.567 | Moderate score; based on four datasets because LUAD was unavailable. |
| 5 | Splatter | 0.600 | Mixed agreement across marginal summaries. |
| 6 | compcodeR | 0.720 | Weaker aggregate marginal agreement than the leading meth-ods. |
| 7 | SimBu | 0.867 | Highest Category A score among the evaluated methods. |

**Table A18:** Top-ranked simulator for Category A within each reference dataset. Scores are dataset-level rank-normalized summaries; lower values indicate stronger agreement.

| **Dataset** | **Samples** | **Top simulator** | **Score** | **Note** |
|---|---|---|---|---|
| TEP | 100 | seqgendiff | 0.000 | Lowest marginal fidelity score in TEP. |
| NSCLC | 25 | SimSeq | 0.167 | Lowest score in the small-sample NSCLC dataset; seqgendiff ranked second. |
| BRCA | 200 | seqgendiff | 0.000 | Lowest marginal fidelity score in BRCA. |
| AD | 150 | seqgendiff | 0.067 | Lowest marginal fidelity score in AD. |
| LUAD | 100 | seqgendiff | 0.040 | Lowest marginal fidelity score in LUAD. |

**Table A19:** Pooled value for the leading simulator on each Category A metric. Values are the unweighted mean of per-dataset means across the five datasets. Lower values indicate closer agreement with the real data.

| **Metric** | **Leading simulator** | **Value** | **Note** |
|---|---|---|---|
| Mean–variance ISE | seqgendiff | 0.1196 | Closest agreement with the real mean–variance trend. |
| logCPM KS | seqgendiff | 0.0139 | Closest agreement with the average logCPM distri-bution. |
| Library-size KS | seqgendiff | 0.0668 | Closest agreement with the sample library-size dis-tribution. |
| Zeros-per-gene KS | seqgendiff | 0.0113 | Closest agreement with gene-level sparsity. |
| Zeros-per-sample KS | seqgendiff | 0.1070 | Closest agreement with sample-level sparsity. |
| BCV KS | seqgendiff | 0.0265 | Closest agreement with dispersion-related BCV sum-maries. |

### Category B: Dependence and multivariate structure

**Table A20:** Category B ranking for dependence and multivariate structure. Scores are rank-normalized category summaries pooled across the benchmark datasets; lower values indicate stronger dependence preservation.

| Rank | Simulator | Category B score | Note |
|---|---|---|---|
| 1 | seqgendiff | 0.292 | Optimal aggregate dependence preservation. |
| 2 | SimSeq | 0.417 | Second-lowest pooled score; strongest for sample-level structure. |
| 3 | Splatter | 0.450 | Competitive pooled score, with favorable behavior in selected datasets. |
| 4 | powsimR | 0.458 | Close to Splatter; based on four-dataset coverage because LUAD was unavailable. |
| 5 | compcodeR | 0.500 | Best on Procrustes alignment, with weaker performance on other dependence summaries. |
| 6 | SPsimSeq | 0.550 | Moderate aggregate dependence preservation. |
| 7 | SimBu | 0.850 | Highest Category B score among the evaluated methods. |

**Table A21:**
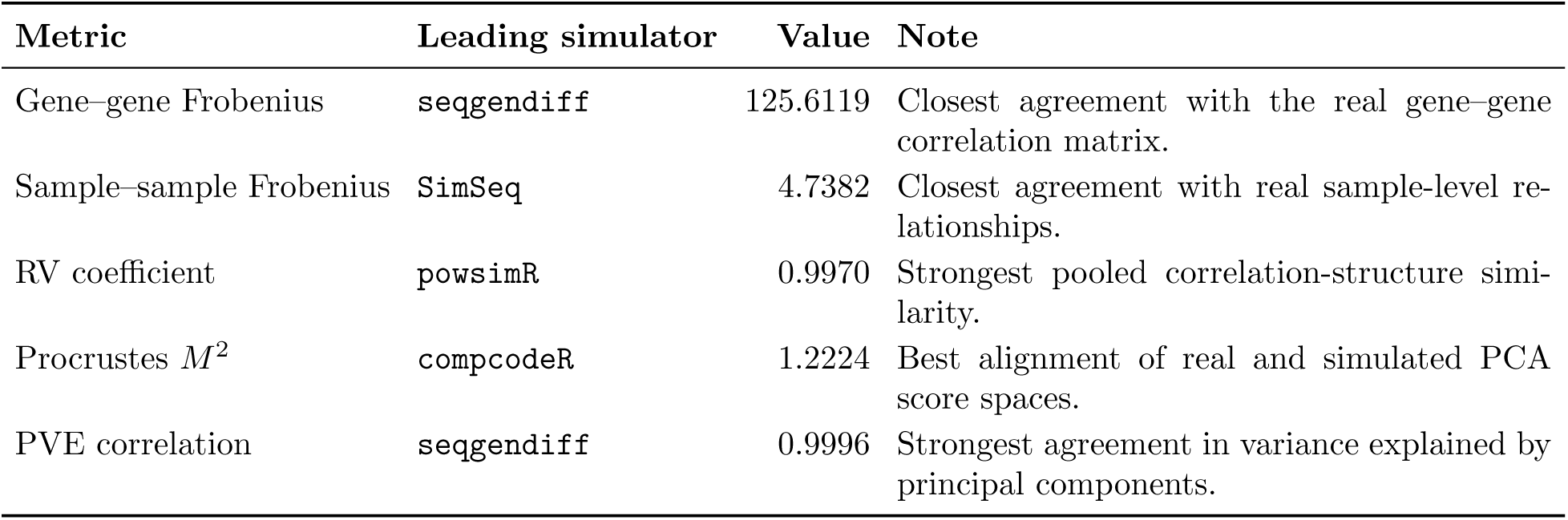
Leading simulator for each Category B metric, pooled across datasets. Lower values are better for the Frobenius distances and Procrustes *M* ^2^; higher values are better for the RV coefficient and PVE correlation.

| **Metric** | **Leading simulator** | **Value** | **Note** |
|---|---|---|---|
| Gene–gene Frobenius | seqgendiff | 125.6119 | Closest agreement with the real gene–gene correlation matrix. |
| Sample–sample Frobenius | SimSeq | 4.7382 | Closest agreement with real sample-level re-lationships. |
| RV coefficient | powsimR | 0.9970 | Strongest pooled correlation-structure simi-larity. |
| Procrustes *M*<sup>2</sup> | compcodeR | 1.2224 | Best alignment of real and simulated PCA score spaces. |
| PVE correlation | seqgendiff | 0.9996 | Strongest agreement in variance explained by principal components. |

**Table A22:** Top-ranked simulator for Category B within each reference dataset. Scores are dataset-level rank-normalized summaries; lower values indicate stronger dependence preservation.

| **Dataset** | **Samples** | **Top simulator** | **Score** | **Note** |
|---|---|---|---|---|
| TEP | 100 | seqgendiff/Splatter | 0.292 | Tied lowest dependence score in TEP. |
| NSCLC | 25 | seqgendiff/Splatter | 0.333 | Tied lowest score in the small-sample NSCLC dataset. |
| BRCA | 200 | compcodeR/seqgendiff | 0.292 | Tied lowest score in BRCA. |
| AD | 150 | SimSeq | 0.208 | Lowest dependence score in AD. |
| LUAD | 100 | seqgendiff | 0.350 | Lowest dependence score in LUAD. |

### Category C: Co-expression network topology

**Table A23:**
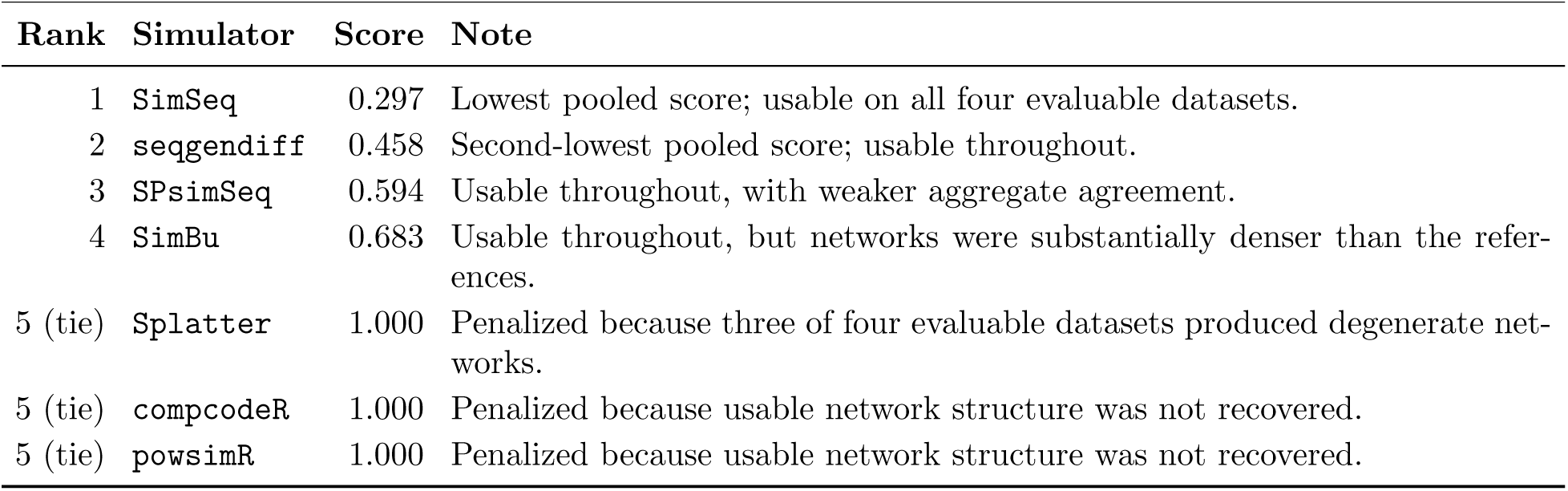
Category C ranking pooled over the four evaluable datasets (BRCA, LUAD, NSCLC, and TEP) after application of the usable-network rule. AD was excluded because the reference network was not evaluable. Lower values indicate stronger agreement with the reference network structure.

| **Rank** | **Simulator** | **Score** | **Note** |
|---|---|---|---|
| 1 | SimSeq | 0.297 | Lowest pooled score; usable on all four evaluable datasets. |
| 2 | seqgendiff | 0.458 | Second-lowest pooled score; usable throughout. |
| 3 | SPsimSeq | 0.594 | Usable throughout, with weaker aggregate agreement. |
| 4 | SimBu | 0.683 | Usable throughout, but networks were substantially denser than the refer-ences. |
| 5 (tie) | Splatter | 1.000 | Penalized because three of four evaluable datasets produced degenerate net-works. |
| 5 (tie) | compcodeR | 1.000 | Penalized because usable network structure was not recovered. |
| 5 (tie) | powsimR | 1.000 | Penalized because usable network structure was not recovered. |

**Table A24:** Leading simulator among the four viable methods for each Category C relative-error metric, pooled across the evaluable datasets. Lower values indicate closer agreement with the real network statistic.

| Metric | Leading simulator | Value | Note |
|---|---|---|---|
| Relative error, density | seqgendiff | 0.7433 | Closest pooled agreement with real network density among the viable methods. |
| Relative error, modularity | seqgendiff | 0.1273 | Closest pooled agreement with real modu-larity. |
| Relative error, average degree | seqgendiff | 0.7433 | Closely tracks density because average de-gree scales with network density. |
| Relative error, clustering | SimSeq / SPsimSeq | 0.1148 | Closest pooled agreement with real local clustering. |

**Table A25:** Top-ranked simulator in Category C for each evaluable reference dataset. Lower scores indicate stronger agreement with the reference co-expression network topology. AD was excluded because the reference network was not evaluable under the benchmark construction.

| Dataset | Samples | Top simulator | Score | Note |
|---|---|---|---|---|
| BRCA | 200 | SimSeq | 0.297 | SimSeq produced the lowest Category C score. |
| LUAD | 100 | seqgendiff | 0.000 | seqgendiff produced the lowest Category C score. |
| NSCLC | 25 | SimSeq | 0.000 | SimSeq produced the lowest Category C score despite the smallest cohort. |
| TEP | 100 | SimSeq | 0.333 | SimSeq produced the lowest Category C score. |
| AD | 150 | - | - | Reference network not evaluable. |

### Category D: Stability across repeated simulations

**Table A26:** Corrected Category D stability scores pooled across the five reference datasets. Lower values indicate less replicate-to-replicate variation. Stability measures reproducibility and should not be interpreted as closeness to the reference data.

| **Rank** | **Simulator** | **Category D score** | **Note** |
|---|---|---|---|
| 1 | compcodeR | 0.133 | Lowest pooled replicate variation. |
| 2 (tie) | SimBu | 0.367 | Relatively low replicate variation. |
| 2 (tie) | Splatter | 0.367 | Relatively low replicate variation. |
| 4 | powsimR | 0.533 | Intermediate replicate variation. |
| 5 | SPsimSeq | 0.600 | Higher replicate variation. |
| 6 | SimSeq | 0.700 | Higher replicate variation. |
| 7 | seqgendiff | 0.800 | Highest pooled replicate variation. |

**Table A27:**
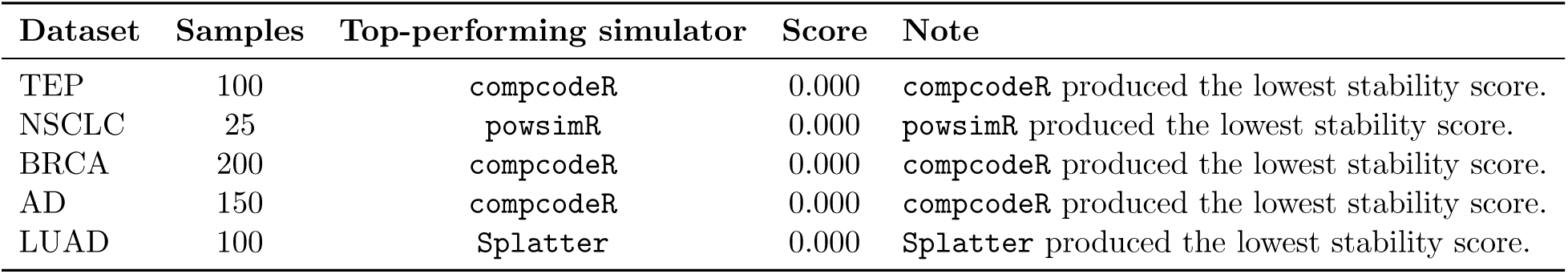
Top-performing simulator in Category D (stability) for each reference dataset. Lower scores indicate greater consistency across the ten independent simulation replicates used for each benchmark dataset.

| **Dataset** | **Samples** | **Top-performing simulator** | **Score** | **Note** |
|---|---|---|---|---|
| TEP | 100 | compcodeR | 0.000 | compcodeR produced the lowest stability score. |
| NSCLC | 25 | powsimR | 0.000 | powsimR produced the lowest stability score. |
| BRCA | 200 | compcodeR | 0.000 | compcodeR produced the lowest stability score. |
| AD | 150 | compcodeR | 0.000 | compcodeR produced the lowest stability score. |
| LUAD | 100 | Splatter | 0.000 | Splatter produced the lowest stability score. |

### Category E: Differential-expression recovery

**Table A28:** Category E ranking for differential-expression recovery, pooled across datasets. Lower values indicate stronger recovery. SimSeq’s score is based on the one dataset where its output could be scored.

| Rank | Simulator | Category E score | Note |
|---|---|---|---|
| 1 (tie) | seqgendiff | 0.167 | Lowest pooled score, with coverage across all five datasets. |
| 1 (tie) | SimSeq | 0.167 | Same pooled score, but based on NSCLC only. |
| 3 | SPsimSeq | 0.250 | Favorable DE recovery, especially in larger cancer datasets. |
| 4 | compcodeR | 0.458 | Moderate pooled score; led several supporting metrics. |
| 5 | powsimR | 0.600 | Weaker pooled DE-recovery score; four-dataset coverage. |
| 6 (tie) | SimBu | 0.900 | high scores and weak DE recovery. |
| 6 (tie) | Splatter | 0.900 | high scores and weak DE recovery. |

**Table A29:** Leading simulator for the primary Category E metrics, pooled across datasets. Higher values indicate stronger recovery.

| Metric | Leading simulator | Value | Note |
|---|---|---|---|
| Recall | SPsimSeq | 0.1135 | Highest pooled recovery of reference DE genes. |
| Jaccard overlap | compcodeR | 0.0583 | Highest pooled overlap of simulated and reference DE gene sets. |

**Table A30:** Top-ranked simulator for Category E within each reference dataset. Lower values indicate stronger DE recovery.

| **Dataset** | **Samples** | **Top simulator** | **Score** | **Note** |
|---|---|---|---|---|
| TEP | 100 | seqgendiff | 0.000 | Lowest DE-recovery score in TEP. |
| NSCLC | 25 | seqgendiff/SimSeq | 0.167 | Tied lowest score in the small-sample NSCLC dataset. |
| BRCA | 200 | SPsimSeq | 0.000 | Lowest DE-recovery score in BRCA. |
| AD | 150 | seqgendiff | 0.000 | Lowest DE-recovery score in AD. |
| LUAD | 100 | SPsimSeq | 0.000 | Lowest DE-recovery score in LUAD. |

**Table A31:** Supporting F1 and ROC AUC highlights for differential-expression recovery. These values were used as supporting summaries rather than as the primary Category E ranking metrics.

| **Dataset** | **Highest-F1 simulator** | **ROC AUC** | **F1 score** | **Note** |
|---|---|---|---|---|
| TEP | seqgendiff | 0.5152 | 0.0532 | Highest F1 in TEP, although recovery re-mained modest. |
| NSCLC | seqgendiff | 0.8051 | 0.2532 | Strongest supporting DE signal among the datasets. |
| BRCA | SPsimSeq | 0.5177 | 0.0758 | Highest F1 in BRCA. |
| AD | SPsimSeq | 0.5058 | 0.0690 | Highest F1 in AD. |
| LUAD | SPsimSeq | 0.4756 | 0.0879 | Highest F1 in LUAD. |

## Supplementary Figures

**Figure A1:**
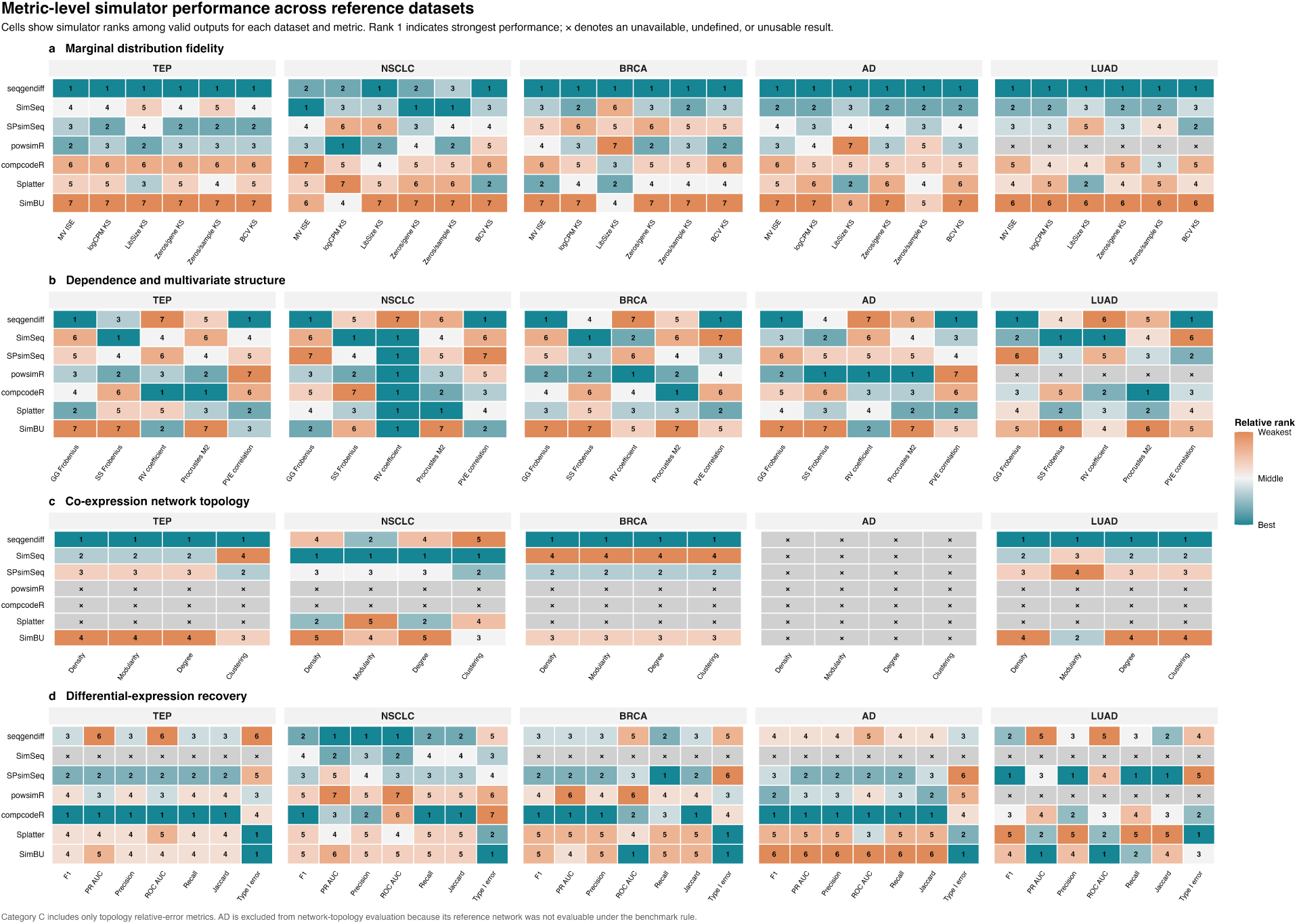
Metric-level simulator ranks across reference datasets for Categories A, B, C and E. Rank 1 denotes the most favourable valid result under the metric-specific scoring direction. A cross denotes an unavailable, undefined or unusable result. Category C shows the topology relative-error metrics, including the supporting average-degree diagnostic, and AD is excluded because its reference network was not evaluable. Category D is shown separately because it is derived from replicate-to-replicate variation.

**Figure A2:**
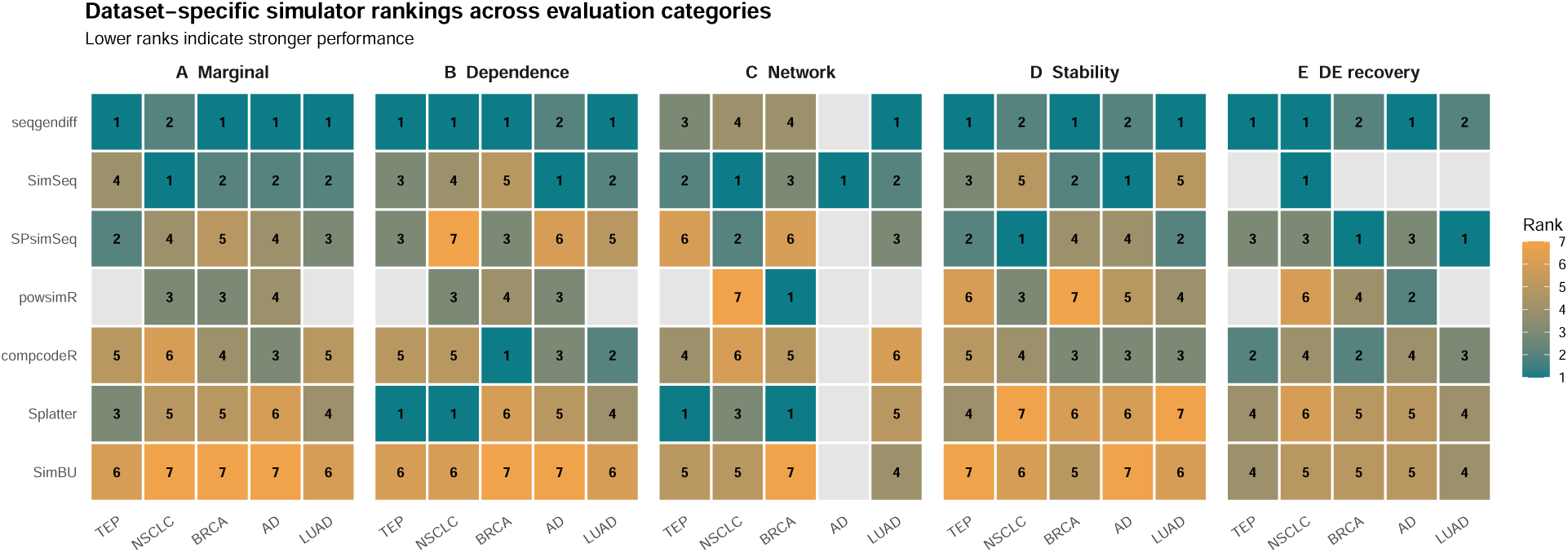
Dataset-specific simulator rankings across evaluation categories. Rankings were computed using all evaluable simulators within each dataset and category. Consequently, LUAD contains six ranked simulators because powsimR was unavailable for LUAD in the fidelity/recovery analyses; Category D was derived separately from the corrected stability analysis.

**Figure A3:**
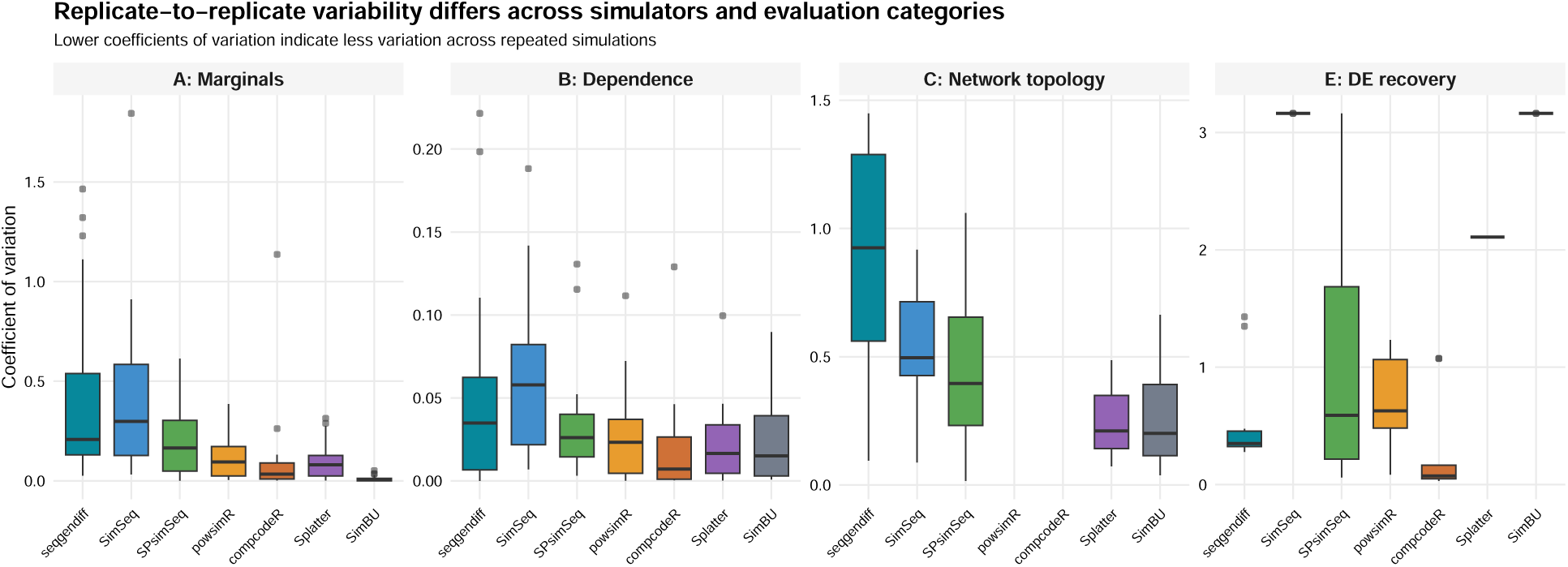
Replicate-to-replicate variation across simulators and evaluation categories. Boxplots show raw coefficients of variation across repeated simulations; lower values indicate less run-to-run variation. These raw coefficients of variation provide the underlying stability measurements from which the normalized Category D benchmark scores were derived.

**Figure A4:**
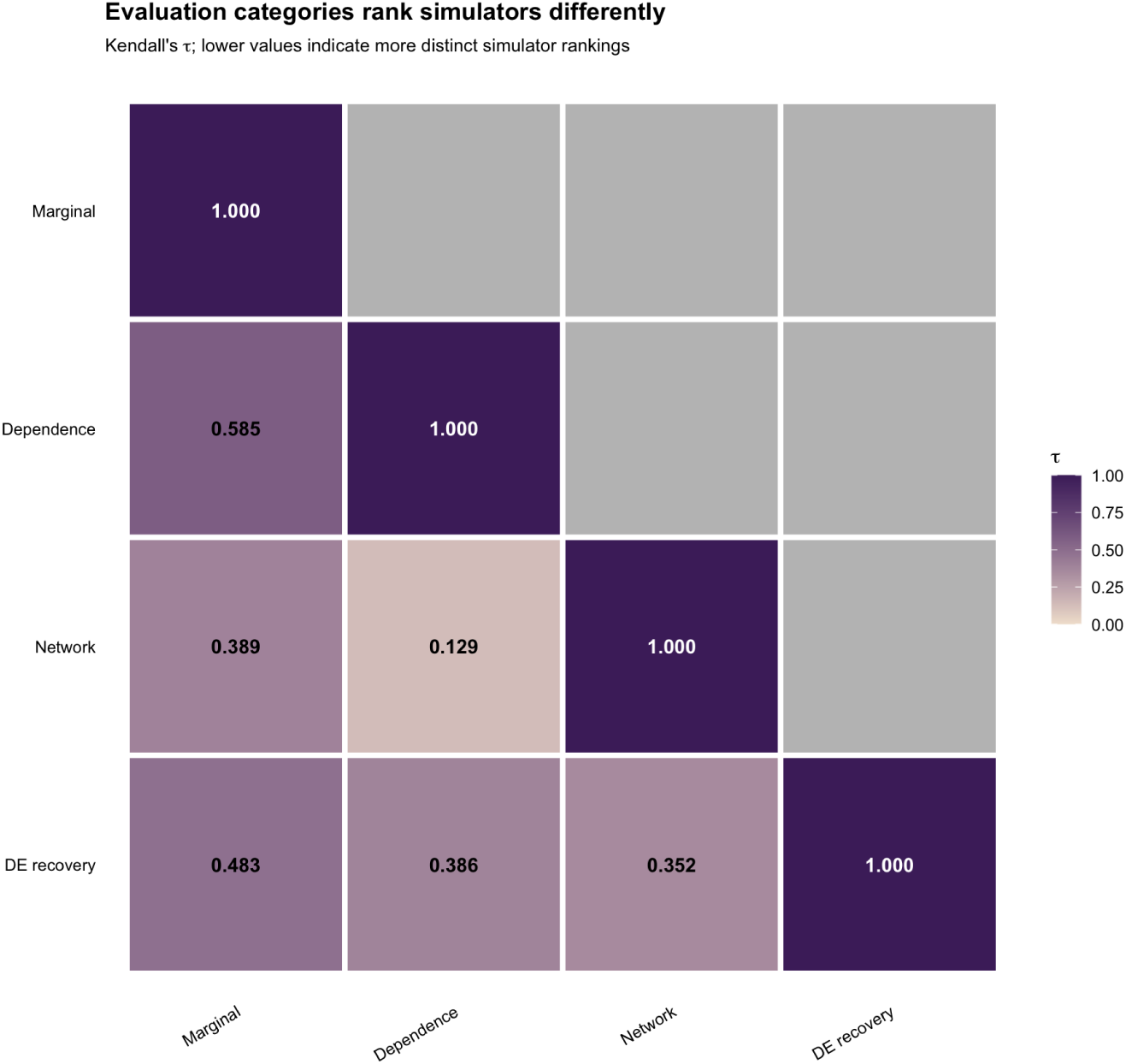
Agreement between simulator rankings across evaluation categories. Cells show Kendall’s rank correlation coefficient *τ* , summarized across datasets for each pair of accuracy-oriented categories. Higher values indicate more similar simulator rankings, whereas lower values indicate greater dissociability between evaluation goals. Category D is excluded because it measures replicate stability rather than agreement with a real-data feature.

**Figure A5:**
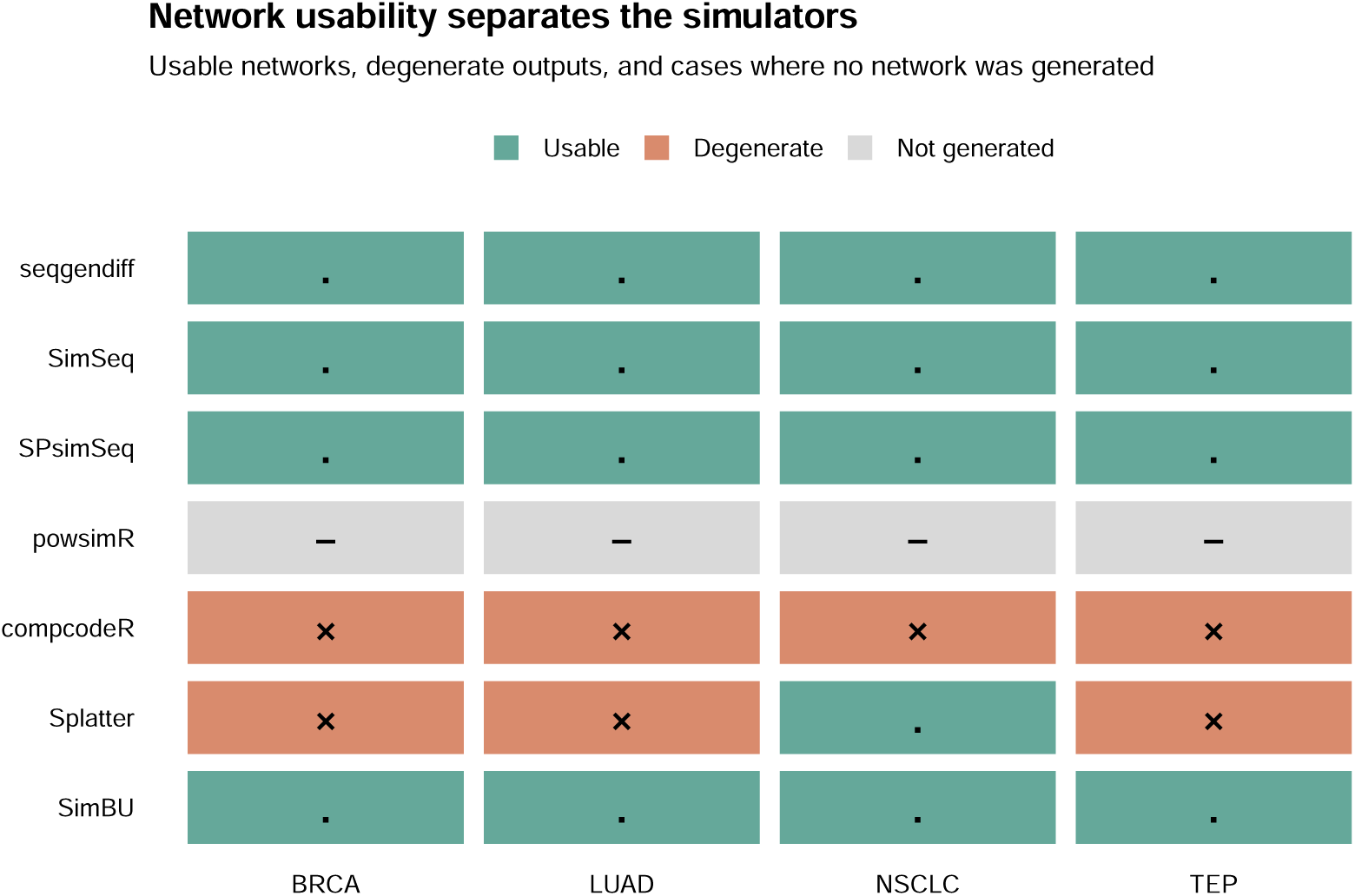
Usability of simulated co-expression networks across the four evaluable reference datasets. Green cells indicate simulator–dataset combinations for which the network contained sufficient structure for Category C topology metrics to be interpreted; orange cells indicate degenerate or otherwise unusable networks under the benchmark rule. AD is not shown because its reference network was not evaluable for Category C. The figure distinguishes failure to generate a usable network from disagreement in topology among networks that were successfully evaluated.

**Figure A6:**
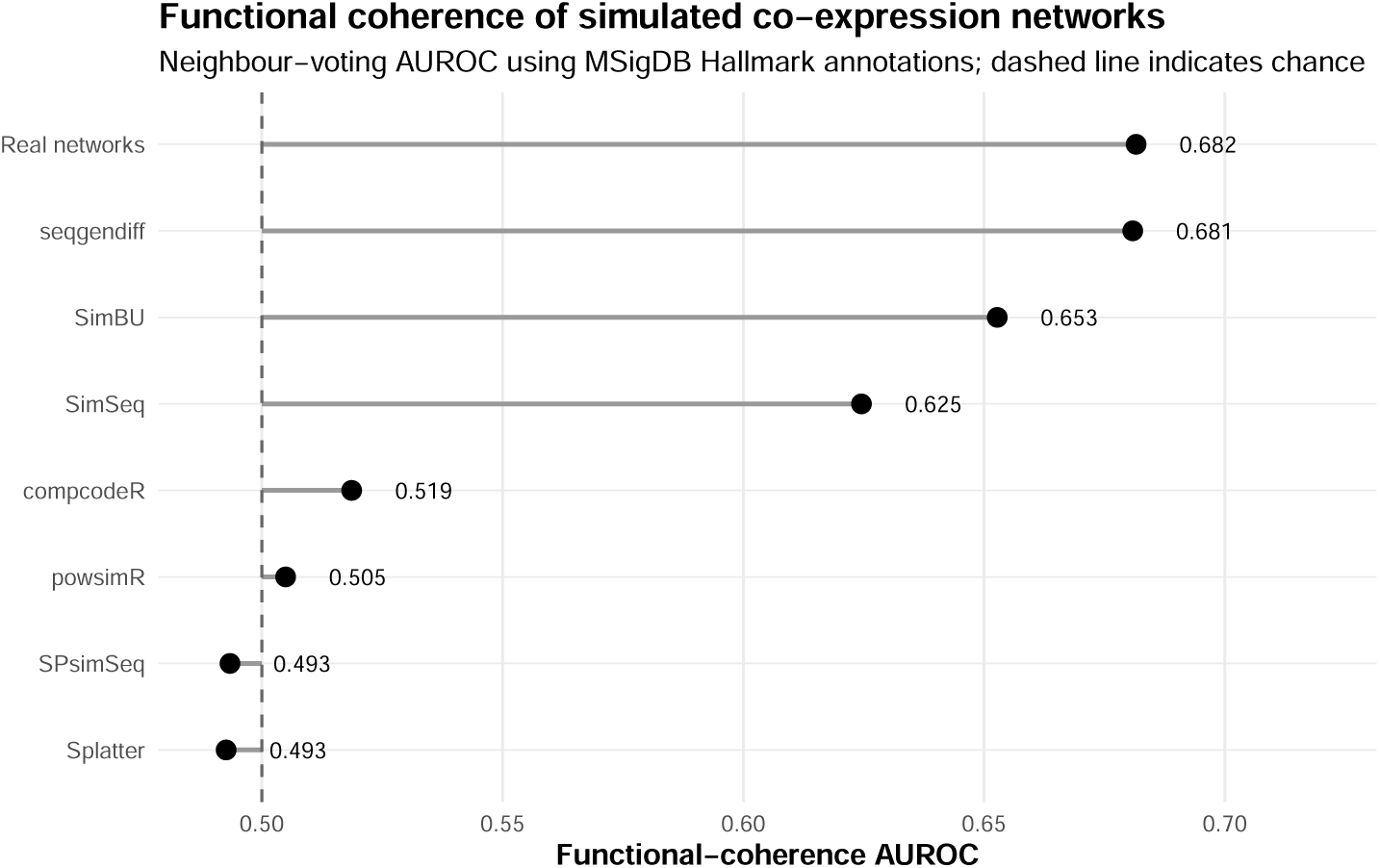
Functional coherence of real and simulated co-expression networks. Points show mean neighbour-voting AUROC across the five reference datasets using MSigDB Hallmark gene-set annotations. The vertical dashed line at 0.5 denotes chance-level prediction. Differences from the corresponding real-network AUROC were calculated within dataset and are summarized in Table A15. This analysis was used as a supporting biological check and was not included in the Category C score.

**Figure A7:**
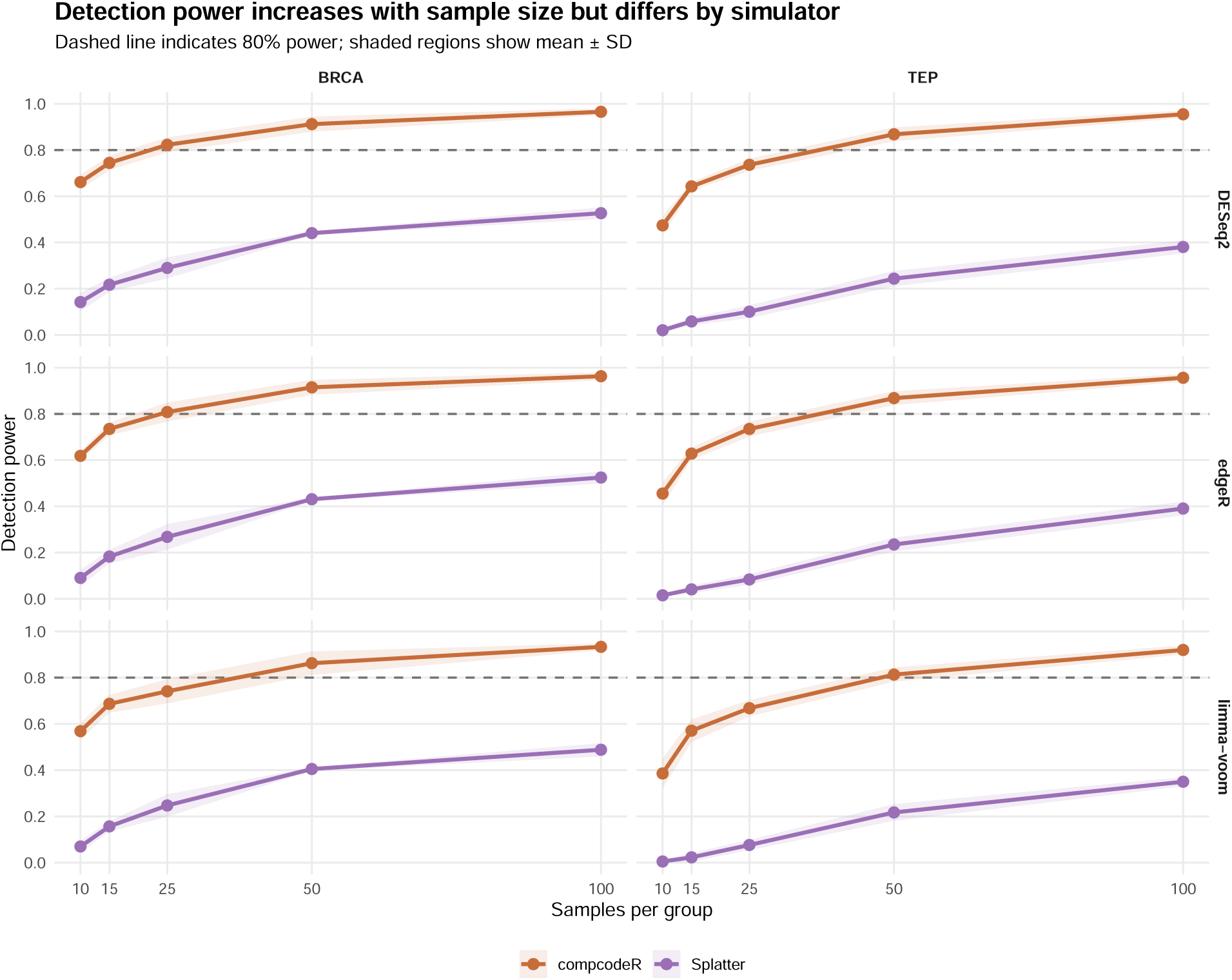
Effect of sample size on differential-expression detection power for compcodeR and Splatter. Curves show mean detection power across increasing numbers of samples per group for BRCA and TEP under DESeq2, edgeR, and limma-voom; shaded regions show mean *±* SD across repeated simulations. The horizontal dashed line marks 80% power. Detection power generally increased with sample size, but its level and rate of increase differed across simulators, datasets, and differential-expression methods, illustrating that simulator choice can influence study-design conclusions.

**Figure A8:**
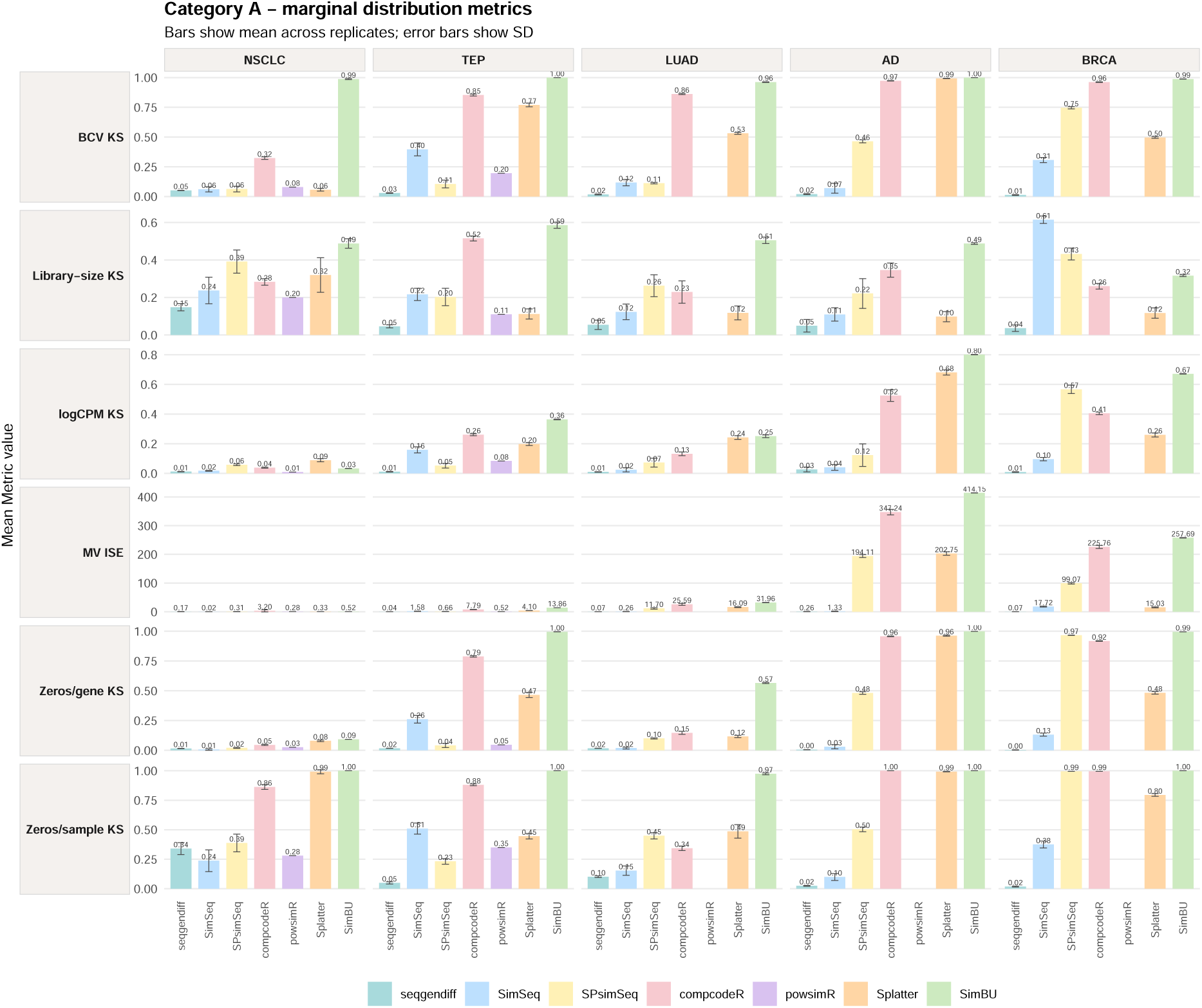
Metric-level comparison of simulator performance for Category A (marginal distribution fidelity) across the five reference datasets. Rows correspond to mean–variance integrated squared error (MV ISE), logCPM distribution, library-size distribution, zeros per gene, zeros per sample, and biological coefficient of variation (BCV). Columns correspond to NSCLC, TEP, LUAD, AD, and BRCA. Bars show the mean metric value across repeated simulations and error bars show the corresponding standard deviation. Metrics are shown on their original scales; lower values indicate closer agreement with the reference data for all Category A metrics. Missing bars indicate unavailable or undefined simulator–dataset results.

**Figure A9:**
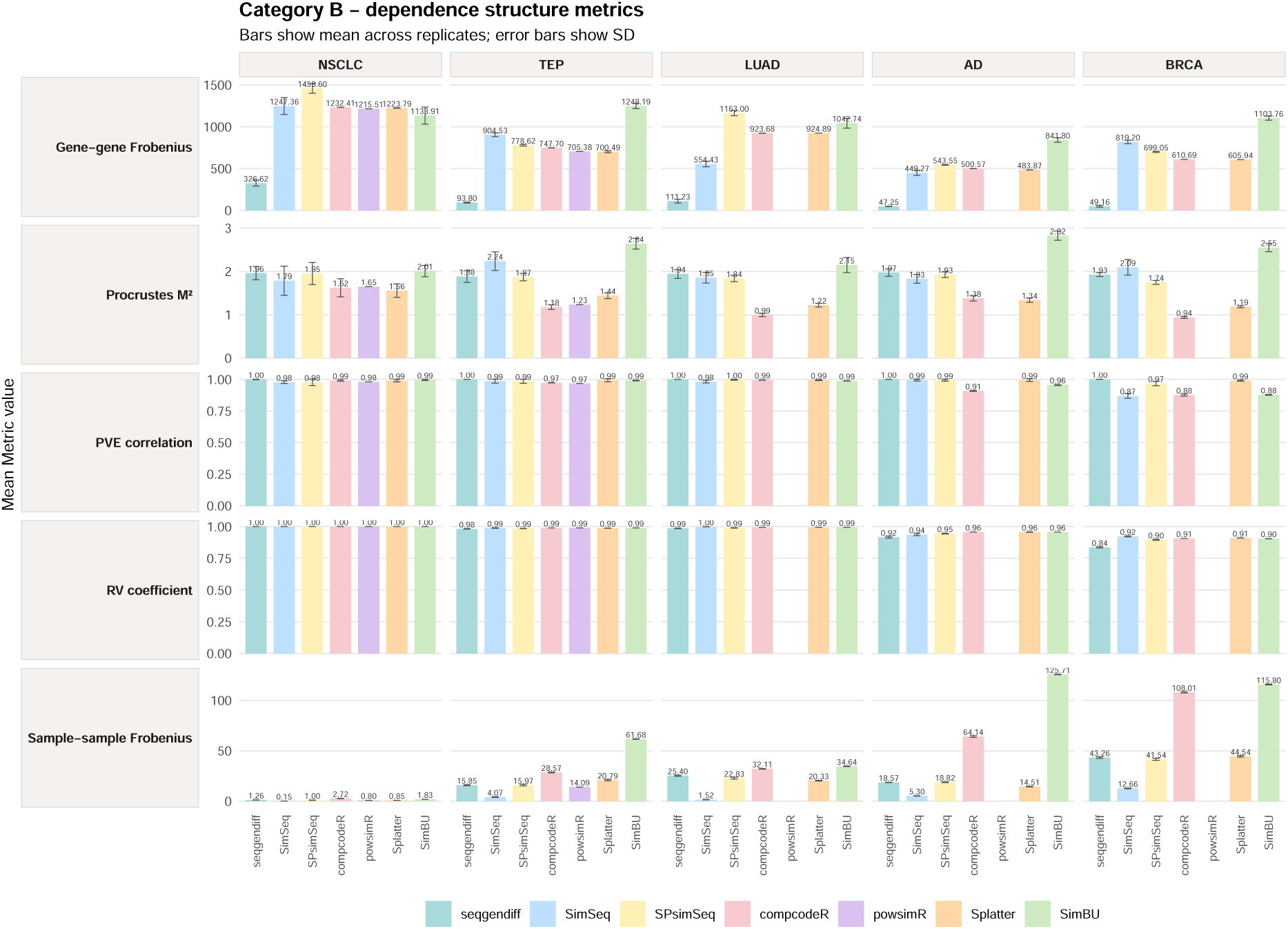
Metric-level comparison of simulator performance for Category B (dependence and multivariate structure) across the five reference datasets. Rows show gene–gene Frobenius distance, sample–sample Frobenius distance, RV coefficient, Procrustes *M* ^2^, and correlation of principal-component variance explained (PVE). Columns correspond to NSCLC, TEP, LUAD, AD, and BRCA. Bars show the mean metric value across repeated simulations and error bars show the corresponding standard deviation. Metrics are displayed on their original scales. Lower values indicate stronger agreement for the Frobenius distances and Procrustes *M* ^2^, whereas higher values indicate stronger agreement for the RV coefficient and PVE correlation. Missing bars indicate unavailable or undefined results.

**Figure A10:**
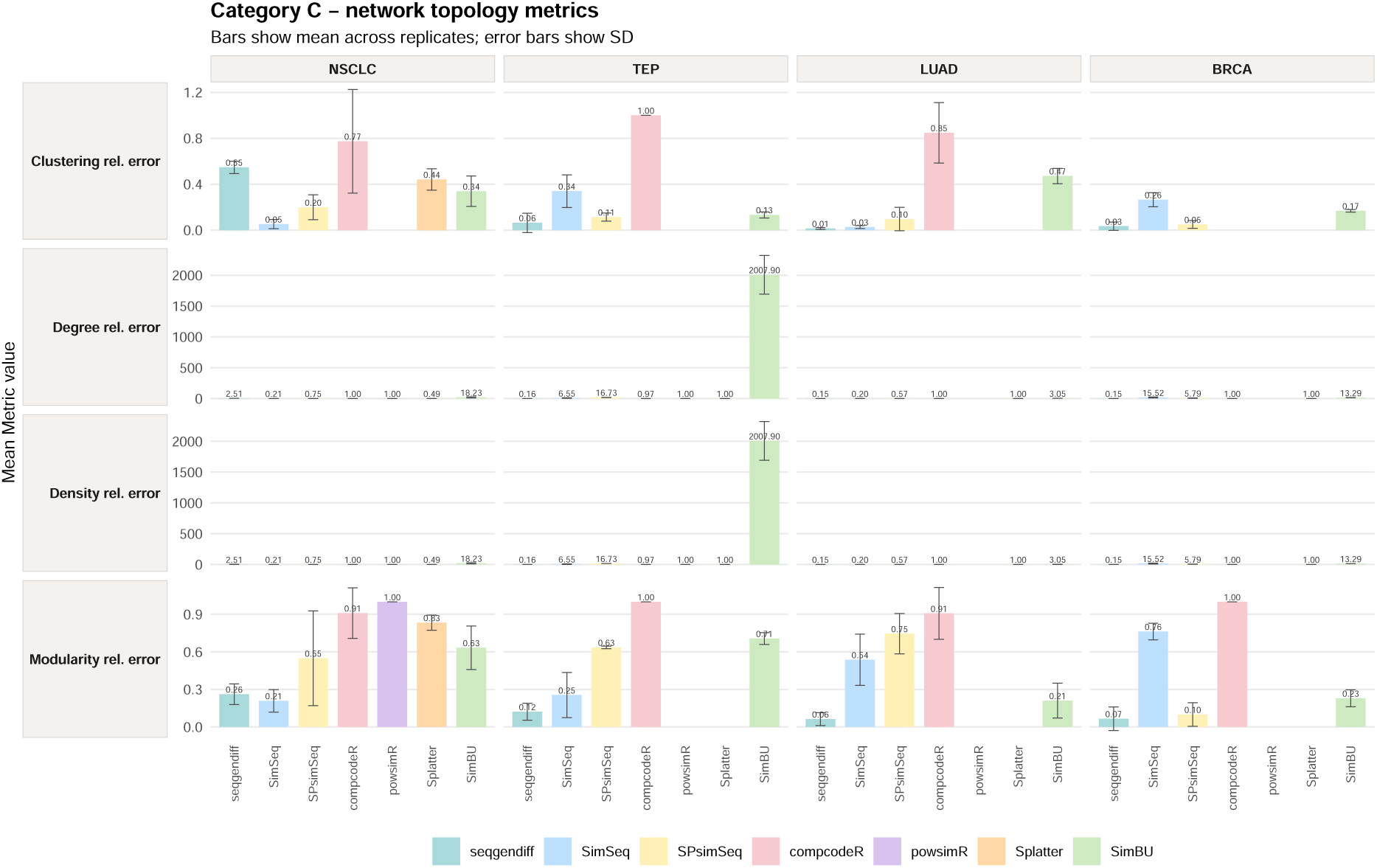
Metric-level comparison of simulator performance for Category C (co-expression network topology) across the evaluable reference datasets. Rows show relative error in network density, modularity, average degree, and clustering coefficient. Bars show the mean metric value across repeated simulations and error bars show the corresponding standard deviation. Lower values indicate closer agreement with the reference network. AD was excluded from the primary Category C analysis because its reference network did not satisfy the predefined network-evaluability criterion. Missing bars indicate unavailable, undefined, or non-evaluable simulator–dataset results.

**Figure A11:**
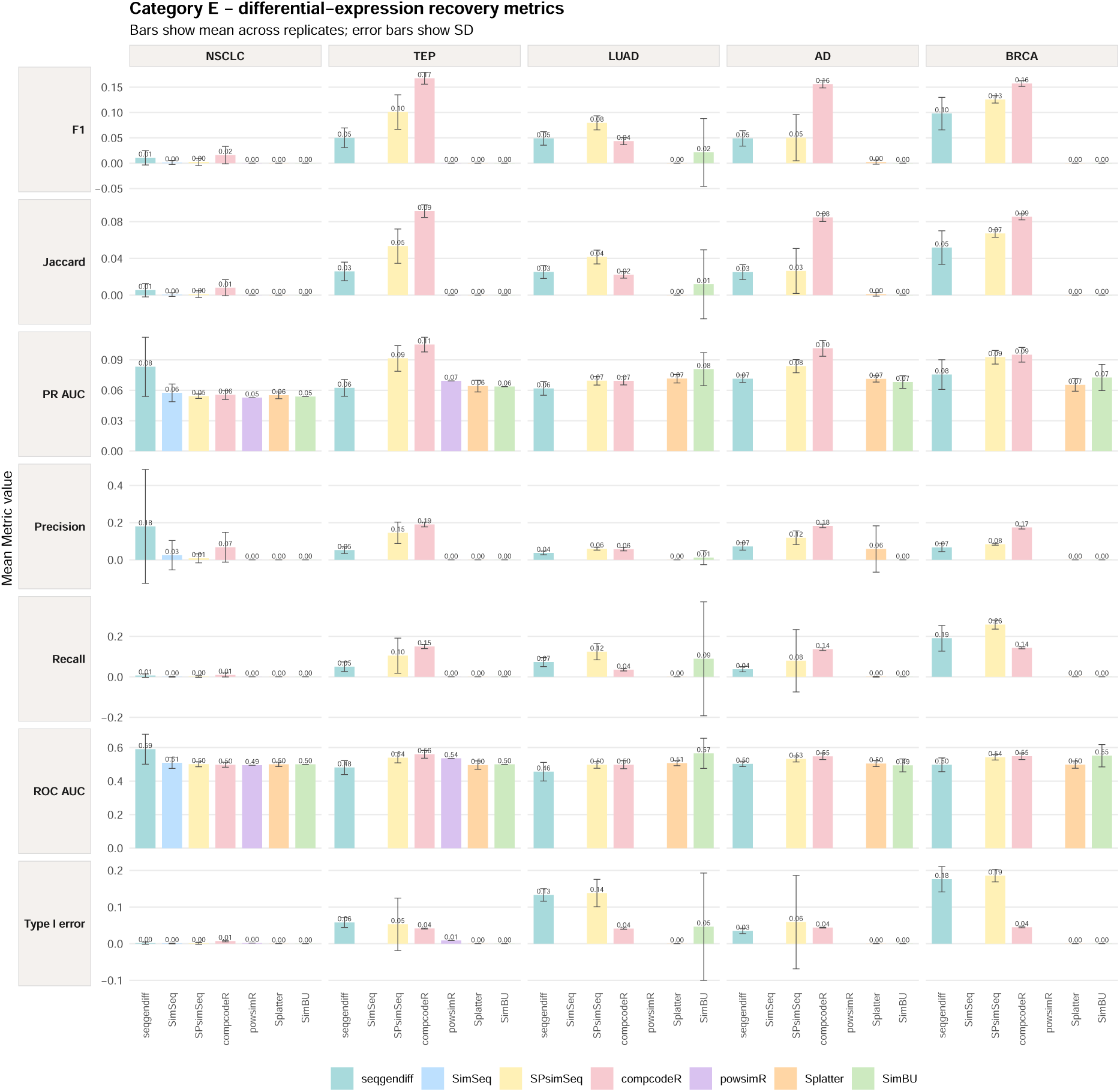
Metric-level comparison of simulator performance for Category E (differential-expression recovery) across the five reference datasets. Rows show F1 score, precision–recall area under the curve (PR AUC), precision, recall, receiver-operating-characteristic area under the curve (ROC AUC), Jaccard overlap, and type I error. Columns correspond to NSCLC, TEP, LUAD, AD, and BRCA. Bars show the mean metric value across repeated simulations and error bars show the corresponding standard deviation. Metrics are displayed on their original scales. Higher values indicate stronger recovery for F1, PR AUC, precision, recall, ROC AUC, and Jaccard overlap, whereas lower values indicate better control of type I error. Missing bars indicate unavailable or undefined differential-expression results.

## Declaration of Generative AI and AI-Assisted Technologies in the Writing Process

During the preparation of this manuscript, the authors used a generative AI assistant for language editing, organization of draft text, and improvement of clarity. The tool was not used to generate data, perform statistical analyses, interpret results, or draw scientific conclusions. All analyses, results, and manuscript content were verified and approved by the authors, who take full responsibility for the work.

## Data Availability

All reference datasets analysed in this study are publicly available from their original repositories and are cited in the manuscript. Processed benchmark inputs, category scores, supplementary result tables, and the files required to reproduce the benchmark analyses will be deposited in a public repository upon publication.

## Code Availability

The code used for data preprocessing, simulator fitting, simulation generation, benchmark evaluation, scoring, statistical analyses, robustness analyses, and figure generation will be deposited in a public repository upon publication.

## Competing Interests

The authors declare no competing interests.

## Notes

### Competing Interest Statement

The authors have declared no competing interest.

### Funding

- National Institutes of Health: P20GM109024

